# Characterization of Rhizosphere Oxidation Associated with Root Development in Rice Using Planar Oxygen Optodes

**DOI:** 10.64898/2026.05.14.725292

**Authors:** Tsubasa Kawai, Shota Teramoto, Xuping Ma, Daichi Fukushima, Kyu Kyu Hmwe, Samuel Munyaka Kimani, Takeshi Tokida, Yusaku Uga

**Affiliations:** Institute of Crop Sciences (NICS), National Agriculture and Food Research Organization (NARO), Tsukuba 305-8518, Japan; Institute for Agro-Environmental Sciences (NIAES), National Agriculture and Food Research Organization (NARO), Tsukuba 305-8604, Japan; Faculty of Biology-Oriented Science and Technology, Kindai University, Kinokawa 649-6493, Japan; Graduate School of Agriculture, Ibaraki University, Ami 300-0393, Japan

## Abstract

Rhizosphere oxidation is a key adaptive mechanism in reductive soil environments, in which oxygen released from roots alters rhizosphere redox conditions and regulates biogeochemical processes. Rice plants possess an internal oxygen transport system, and radial oxygen loss (ROL) from roots is closely associated with root development. However, the spatial patterns of ROL in soil and their relationships with root traits remain poorly characterized. In this study, we developed a multimodal imaging system that integrates planar oxygen optodes with X-ray computed tomography to simultaneously visualize rhizosphere oxidation and root development in rice. Daily time-course tracking of individual crown roots revealed dynamic changes in the spatial distribution and magnitude of rhizosphere oxygen in relation to root elongation and aging. Root thickness was positively correlated with dissolved oxygen levels near root tips. Genotypic comparisons further identified a cultivar with reduced rhizosphere oxidation despite possessing thicker roots among the tested genotypes, thereby indicating the involvement of additional physiological processes. Overall, these findings demonstrate that rhizosphere oxidation is regulated by root growth stage and thickness and dynamically modulated during root development.

## Introduction

Among crop species, rice (*Oryza sativa* L.) is uniquely adapted to flooded soil environments because of its ability to transport and release oxygen from its roots. Rice is predominantly cultivated in flooded paddy fields, where persistent waterlogging rapidly depletes soil oxygen levels. Under these deoxygenated conditions, anaerobic microbial processes promote the accumulation of various phytotoxic compounds, including ferrous iron (Fe^2+^) and reduced sulfur species (Drew and Lynch, 1980). To mitigate these effects, rice roots generate a protective oxygenated zone via oxygen leakage from root tissues, a process known as radial oxygen loss (ROL) (Colmer, 2003a). Rhizosphere oxidation serves as a key adaptive mechanism that detoxifies reduced toxic compounds and supports plant survival in strongly reductive soil environments.

Rice maintains an efficient internal oxygen transport system that delivers oxygen from the shoot to respirating root tissues, particularly at the root tips. Lysigenous aerenchyma—gas-filled spaces formed through cortical cell death and lysis—provides a continuous pathway from the leaves to the stems and into the roots (Colmer and Pedersen, 2008; Yin et al., 2021). Oxygen diffuses longitudinally through the aerenchyma along concentration gradients determined by oxygen consumption during root respiration and ROL (Pedersen et al., 2009). ROL barrier formation in the outer cell layers limits radial oxygen diffusion in the basal regions and facilitates efficient oxygen transport to respiring root tips, leading to the predominance of ROL at the root tips rather than in the basal regions (Colmer et al., 1998; Ejiri et al., 2021). This barrier is largely associated with suberin deposition in the exodermis (Shiono and Nakazono, 2024), and recent evidence indicates that glycerol ester accumulation contributes to its tightness (Jiménez et al., 2024a, 2025). Therefore, aerenchyma formation, ROL barrier development, and respiratory oxygen consumption determine the spatial extent and magnitude of ROL.

The ROL extent and spatial distribution are strongly influenced by root developmental stage and structural traits. ROL barriers typically form in longer, mature roots but are less common in shorter roots (< 40 mm) under stagnant solution conditions (Colmer et al., 2006; Ejiri et al., 2021). In longer roots, aerenchyma formation begins in relatively young regions near the root apex (10–20 mm from the root tips) and gradually extends toward the basal region. Conversely, ROL barrier formation is initiated in more mature tissues, typically beyond 20–30 mm from the root tips (Kotula et al., 2009; Yamauchi et al., 2018). This temporal and spatial gap generates a distinct ROL-active zone in the apical parts of the roots. Root thickness further influences the extent of ROL near the root tip because thicker roots generally develop a larger aerenchyma area and consequently release more oxygen at their tips (Yamauchi et al. 2019). However, these developmental and structural relationships have been primarily observed under stagnant solution conditions, and their relevance to rhizosphere oxidation patterns in soil environments remains unclear.

Genotypic variations in the spatiotemporal patterns of ROL under stagnant solution conditions have been previously reported in rice. Colmer (2003b) demonstrated that five lowland and deep-water rice genotypes formed tight ROL barriers, restricting ROL from basal root regions while maintaining variable ROL levels at the root tips. In contrast, two of seven upland genotypes did not form ROL barriers and displayed detectable ROL along basal regions (Colmer, 2003b). Similarly, another upland genotype was reported to lack an ROL barrier (Colmer et al., 1998), indicating evolutionary divergence in ROL-related traits among rice ecotypes. Several studies have indicated that aerenchyma formation rate is involved in oxygen release from roots at the whole-root-system level across genotypes (Zheng et al., 2018; Chen et al., 2019; Iqbal et al., 2021; Wei et al., 2025). However, the relationship between genotypic variation in root anatomical traits and the spatiotemporal patterns of ROL during root development in rice remains inadequately understood.

ROL from rice roots has traditionally been quantified using single-point measurements of dissolved oxygen (DO) concentrations with cylindrical O_2_ cathodes or microelectrodes (Jiménez et al. 2021). Measurements at multiple positions along the root axis using cylindrical O_2_ cathodes have been used to characterize longitudinal ROL profiles in hydroponically cultivated roots (Armstrong and Wright, 1975). Microelectrodes provide high-resolution oxygen measurements by penetrating tissues, enabling micrometer-scale profiling both outside and inside the roots (Armstrong, 2000). While these methods have advanced understanding of ROL distribution and associated plant traits, they are largely limited to hydroponic systems, leaving soil oxygen dynamics poorly understood. Planar optodes provide a non-destructive, two-dimensional luminescent imaging approach for visualizing rhizosphere dynamics of O_2_, pH, pCO_2_, and ammonia (Blossfeld, 2013). They offer spatial and temporal resolutions comparable to those of microelectrodes while also enabling visualization across larger regions of interest (Larsen et al., 2011). Larsen et al. (2015) used planar optodes in anoxic soils to demonstrate dynamic changes in oxygen distribution in the rice rhizosphere. They showed that ambient conditions, including light–dark transitions and irradiance, influenced the extent of the oxic zone around the roots. Spatiotemporal variations associated with root emergence, aging, and lateral root formation have also been observed (Larsen et al., 2015). Shiono et al. (2022) reported dynamic changes in oxygen status during rice seed germination as the coleoptile elongated and emerged from anoxic conditions in the atmosphere. Combining planar optodes with advanced imaging techniques has further enabled investigation of the relationships between ROL and rhizosphere ion dynamics, including those of iron (Williams et al., 2014; Maisch et al., 2019, 2020), phosphate (Fang et al., 2021), and organic carbon (Wei et al., 2025), in rice. These findings demonstrate the utility of planar optodes for examining the spatiotemporal dynamics of rhizosphere oxidation. However, the spatiotemporal patterns of ROL along the root axis during root elongation and their relationships with root traits remain poorly characterized in soil environments, despite being well established under stagnant solution conditions.

In this study, we aimed to develop a multimodal imaging system that integrates planar oxygen optodes with X-ray computed tomography (CT) to simultaneously visualize rhizosphere oxidation and root development in rice. X-ray CT enables non-destructive visualization of root system architecture (RSA) in soils based on differences in X-ray attenuation between root tissues and soil particles (Heeraman et al., 1997; Teramoto et al., 2020). By combining these techniques, we attempted to elucidate how rhizosphere oxidation dynamically changes with root growth under anoxic and highly reduced soil conditions. Furthermore, we examined genotypic differences in rhizosphere oxidation patterns to identify the root traits associated with this process.

## Materials and methods

### Plant materials and field observations

Four lowland rice (*Oryza sativa* L.) cultivars were used: ‘Koshihikari’ (KSH, *japonica*), ‘Muha’ (*aus*), ‘Qiu Zao Zhong’ (QZZ, *indica*), and ‘Tupa121-3’ (*aus*). KSH is widely cultivated in Japan, whereas the other cultivars were selected from the world rice core collection and reported to exhibit diverse root morphology and anatomy (Uga et al., 2009). Field observations of root systems for these cultivars were conducted at a paddy field of the National Agriculture and Food Research Organization (NARO) in Tsukubamirai, Japan (36°00′ N, 140°01′ E) in 2022. Crop management and field conditions followed those described by Hamamoto et al. (2025). Root sampling was conducted using the pinboard method, modified from Uga et al. (2013), at three distinct developmental stages (panicle initiation, booting, and grain filling). After soil removal, root systems were photographed. The resulting images revealed cultivar- and stage-dependent differences in root volume and coloration (Fig. S1), highlighting morphological and physiological variation under reductive field conditions.

### Rootbox preparation and plant cultivation

An oxygen-sensitive optode foil (100 mm width × 75 mm height; SF-RPSu4; PreSens GmbH, Regensburg, Germany) was attached to a glass plate (108 mm width × 198 mm height × 2 mm thickness; TEMPAX Float^®^; AS ONE, Osaka, Japan). The glass plate was subsequently affixed to an open-top polystyrene box (108 mm width × 194 mm height × 18 mm thickness; AS ONE) with the optode foil oriented toward the interior of the rootbox (Fig. S2A, B). The gaps between the glass plate and box were sealed using silicone sealant.

Soil was collected from the paddy field at the NARO in Tsukubamirai, Japan. The soil was classified as clay loam (USDA classification). It was crushed, sieved through a 2-mm mesh, and air-dried in a greenhouse environment. Prior to use, the dried soil was thoroughly mixed with water and poured into the rootbox. The soil surface was adjusted to approximately 15 mm above the upper edge of the optode foil.

Seeds were surface-sterilized using a fungicide (Plant Preservative Mixture; Plant Cell Technology, Washington, D.C., USA) at 15 °C for 24 h and pre-germinated in water at 30 °C for 2 days. Subsequently, seeds were placed on floating mesh in one-quarter-strength Kimura B solution (Yoshida et al., 1976) under long-day conditions (14 h light/10 h dark) with a graded temperature regimen (20–30 °C), 50–75% relative humidity, and a light intensity of 400–470 μmol m^-2^ s^-1^ at 400 ppm CO_2_ in a growth chamber (LPH-411PFQDT-SPC; Nippon Medical & Chemical Instruments Co., Osaka, Japan) (Fig. S3A–D).

One day after soil filling, 7-day-old seedlings were transplanted into the center of the soil surface at a depth of 5–10 mm. Following transplantation, light intensity was increased to 400–700 μmol m^-2^ s^-1^ (Fig. S3D). Four rootboxes were placed in a 6-L square bucket (230 mm × 130 mm × 250 mm) at a 60° angle, with the optode foil on the back. Soil surfaces were covered with black plastic sheets to inhibit light penetration (Fig. S2C). The bucket was filled with water, and the soil in each rootbox was maintained under flooded conditions throughout cultivation. For each cultivar, two biological replicates were grown. The experiment was repeated twice, resulting in four replicates analyzed per cultivar.

### Analysis of dissolved Fe^2+^ concentration in the rootbox

Soil reductive status in the rootbox, used for shoot and root morphological analyses, was assessed by collecting soil solution from porous cups (φ3 mm × 20 mm) vertically inserted adjacent to each plant at 40–60 mm depth. The soil solution was sampled 20 days after sowing (DAS) (Kajiura et al., 2025). Fe^2+^ concentrations in the collected solution were measured using inductively coupled plasma optical emission spectrometry (ICP-OES 720-ES; Agilent Technologies, CA, USA). The soil solution collected from the rootboxes without plants served as the blank control.

### Planar optode imaging of oxygen

A flowchart that summarizes the data collection and analysis processes is presented in Figure S4. Oxygen distribution in the rhizosphere was analyzed using a planar optode imaging system (VisiSens TD; PreSens GmbH; Koop-Jakobsen et al., 2018). The imaging platform was set up in a dark room and equipped with a camera linked to a PC running the imaging software (VisiSens ScientifiCal version 1.0.2.9, PreSens GmbH), an light-emitting diode (LED) light source (VisiSens TD Big Area Imaging Kit, PreSens GmbH), and a custom-made rootbox stage with LED lights for plant illumination (PF10-S4WT8-D; Nippon Medical & Chemical Instruments Co.) (Fig. S2D–F). Both the camera, mounted on a custom-made XYZ stage (AS ONE), and rootbox stage were placed on an aluminum breadboard (300 mm × 900 mm; Edmund Optics, NJ, USA) inclined at 15°.

In this system, oxygen concentration was determined using a ratiometric approach (Larsen et al., 2011). The camera captured the fluorescence emitted from the optode foil, which contained O_2_-sensitive indicator (red fluorescence) and stable reference (green fluorescence) dyes, both excited by the LED light source. The red fluorescence of the indicator dye was quenched by O_2_ molecules; therefore, the red-to-green (R/G) fluorescence ratio reflected the O_2_ concentration.

For each measurement, the rootboxes were gently placed on the imaging platform, while optode images were recorded. The exposure time was set to 2.5 ms, with light intensity at its maximum. The spatial resolution was approximately 105 μm per pixel (9.5 pixels per mm). Imaging started one week after transplantation and continued for two weeks, with measurements conducted daily between 10:00 and 14:00 (Fig. S3A–D).

### Optode foil calibration

Calibration was performed following the final measurement, as previously described (Shiono et al., 2022). Rootboxes were initially filled with water and deoxygenated by flushing with N_2_ gas until the DO value reached <0.1 mg L^-1^. At each stage, optode images were captured under the same conditions used for the measurements (30 °C), while simultaneously recording the DO value and temperature with a DO meter (FireSting^®^-GO2, OXROB10-OI, and TSUB21; PyroScience GmbH; Aachen, Germany). Subsequently, the water was gradually oxygenated by flushing it with air, and the optode images and DO values were recorded at discrete stepwise oxygenation levels. Finally, sodium dithionate [Na_2_S_2_O_4_; final concentration 0.16% (g v^-1^); FUJIFILM Wako Pure Chemical Corporation, Osaka, Japan] was added to the water to obtain an anoxic image. In total, approximately 15 calibration points covering 0–100% air saturation were collected. The average R/G ratio for each optode image was obtained using an imaging software (VisiSens ScientifiCal; PreSens GmbH). The DO values were converted to air-saturation percentages, and calibration curves were generated in Microsoft Excel 365. The calibration curve remained consistent before and after the cultivation and imaging measurements (Fig. S5).

### Imaging of root structure by X-ray computed tomography

The opaque nature of the optode foils, which absorbs excitation light and thereby prevents autofluorescence from roots and soil behind the optode from interfering with the measurements, necessitated the use of X-ray CT to visualize RSA. Following the optode measurements, four rootboxes were placed in a 6-L square bucket, and CT scanning was performed using an X-ray CT system (Fig. S2G, H; inspeXio SMX-225CT FPD HR; Shimadzu Corporation, Kyoto, Japan). The CT scanning conditions were as follows: tube voltage, 225 kV; tube current, 500 μA; copper filter, 0.5 mm thick; and total scanning time, 10 min. A total of 792 tomography images were reconstructed into 3D volumes (1,024 × 1,024 pixels; 16-bit), with 199-μm voxel resolution.

The 3D image volumes were processed using VGSTUDIO MAX software (version 3.2.5.160436; Volume Graphics GmbH, Heidelberg, Germany), and 2D root system images corresponding to 1-mm-thick slabs immediately below the optode foils were extracted. Root diameter was measured at positions 1–3 cm from the basal region using ImageJ software (https://imagej.net/ij/).

### Image processing and dissolved oxygen mapping

All optode and CT images were aligned using GIMP software (version 3.0.4; https://www.gimp.org/). Optical distortion in optode images was corrected by applying the “Lens Distortion” filter in GIMP with the main parameter set to −15. The CT images were aligned with the optode images to maintain spatial resolution. The aligned optode and CT images were exported as PNG files at the same dimensions.

Based on the time-course images, crown roots that elongated along the optode foil and maintained contact with the foil for a minimum of two consecutive days were selected for DO profile analysis. A Python-based program, “RG2DO-Root”, was developed to extract DO values along root axes. The CT image from the final measurement and merged time-course optode images were imported, allowing for the visualization of the DO distribution by overlaying optode-derived DO values onto the CT image. The root axis of the selected crown roots was manually traced from the stem base by sequentially clicking along the root path in the CT image. R and G intensity values (0–255) were extracted for each pixel along the traced line in all optode images. The R/G ratio was calculated and converted into DO values using a calibration curve obtained after cultivation (Fig. S5). At each point along the traced line, the average DO value was calculated within a circular region centered on the point with a radius of 0–10 pixels (Fig. S6). In addition, the number of pixels with DO >10% within the designated circular area was determined, and their proportion to the total number of pixels in the circle was calculated.

The measurement noise boundary was evaluated by analyzing the calibration images acquired at six DO levels ranging from 0 to 20% air saturation, corresponding to the range observed in the rhizosphere measurements. The DO value distributions were measured along the central vertical line using RG2DO-Root with integration radii of 0 and 10 px (Fig. S7A–D). In a homogenous anoxic solution, the mean DO value measured with RG2DO-Root was 1.4%, while the upper noise boundary, defined as the 95th percentile, was 4.3% when the integration radius was set to 0 px (point-wise measurement). In contrast, with an increase in the integration radius to 10 px, the mean DO value remained similar (1.3%), whereas the upper noise boundary substantially decreased to 1.9%. This noise reduction caused by increasing the measurement area was consistent across all calibration DO levels (Fig. S7E, F).

### Extraction of dissolved oxygen profiles along the crown root axis

DO values along selected crown roots were extracted from the DO matrix derived from optode measurements. Daily root elongation was quantified using ImageJ software. DO profiles were integrated with daily root-length data to track oxygen dynamics along elongating crown roots using the R packages dplyr and tidyr in R (version 4.2.1; R Core Team, 2022). At each defined distance from the root tip, the DO value of the nearest pixel was selected for quantification. Background DO was defined as the minimum DO value at the corresponding spatial position over the preceding three days. For the continuous quantification of DO along a crown root, DO values were extracted along a line shifted 20 pixels to the right of the root axis and used as background DO.

### Analyses of shoot and root morphological traits

The four rice cultivars were cultivated in open-top polystyrene boxes with lids (194 mm × 108 mm × 26 mm; AS ONE). Pre-germinated seeds were sown in paddy soil at a density of two seeds per box. Following seedling emergence, the water level was increased and maintained to ensure continuous flooding. All other growth conditions were identical to those employed for the optode experiments.

At 21 DAS, the leaf age, shoot length, and tiller number were recorded. Rootboxes were imaged using X-ray CT, while root system images were extracted using the same parameters as those used for optode measurements to visualize the root system. The root cone angle was quantified from processed CT images using ImageJ software.

After CT scanning, roots were carefully washed to remove soil and scanned using a flatbed scanner at 400 dpi (Expression 12000XL; Seiko Epson Corporation, Nagoya, Japan). Crown roots were counted manually. Root images were analyzed using WinRHIZO Pro 2017a software (Regent Instruments, Quebec, Canada). Root length was classified into two diameter classes (<300 µm, lateral roots; ≥300 µm, crown roots). Shoots and roots were oven-dried at 80 °C for more than three days to determine shoot and root dry weights. Root tissue density was calculated by dividing root dry weight by root volume measured using WinRHIZO.

### Analysis of root anatomical traits

The four rice cultivars were cultivated under the same conditions as those described for morphological analysis. At 21 DAS, root segments located 1–2 cm from the base were sampled, and root diameter was measured using a microscope (Panthera E2; Shimadzu, Kyoto, Japan) equipped with a micrometer.

For anatomical analysis, elongating crown roots approximately 10 cm in length were selected. These roots exhibited lateral roots exclusively in the basal region, with the lateral root formation zone extending less than half that of the crown root length. Root segments were sampled at distances of 8–12, 18–22, 28–32, 38–42, and 48–52 mm from the root tip and subsequently fixed in FAA solution [5% formaldehyde, 5% acetic acid, 63% ethanol (v v^-1^)]. Samples were deaerated under vacuum for 30 min. The FAA solution was then replaced, and samples were stored at 4 °C. Prior to sectioning, root samples were rinsed with water and embedded in 3% agarose. Roots were sectioned into 100-µm-thick slices using a microslicer (DTK-1000; Dosaka, Kyoto, Japan) and examined under a microscope (AX70; Olympus, Tokyo, Japan). Tissue layer thickness, total cross-sectional area, and aerenchyma area were quantified using ImageJ software. The proportion of aerenchyma was calculated by dividing the aerenchyma area by the cortical area.

Suberin lamella formation was analyzed using a rapid staining method (Yamauchi et al., 2025). The root segments were deaerated in water under vacuum for 30 min. The samples were embedded in 3% agarose and sectioned into 100-µm-thick slices. Agarose was removed by heating the sections at 80 °C for 10 min. Suberin lamellae were stained with 0.01% fluorol yellow 008 (FY088; g v^-1^ in 99.5% EtOH) at 60 °C for 10 min. Subsequent to staining, the solution was removed and sections were rinsed with distilled water. FY088 fluorescence was observed using a laser-scanning microscope (LSM710; Carl Zeiss, Oberkochen, Germany) with excitation set at 488 nm and detection at 520–580 nm. Transmitted light images were acquired using a T-PMT detector.

### Statistical analysis

Differences in morphological and anatomical characteristics, DO concentrations, and soil chemical conditions were analyzed using two-tailed Student’s *t*-test (for pairwise comparisons) or one-way ANOVA followed by Tukey’s multiple-comparison tests, implemented with the glht function in the multcomp package (version 1.4-20; Hothorn *et al*., 2008) in R. Pearson’s correlation analysis was conducted using the corrplot package (version 0.92; Wei and Simko, 2021).

## Results

### Development and validation of a multimodal imaging workflow for rice rhizosphere oxidation

To link rhizosphere oxygen dynamics to root development in soil, a multimodal imaging workflow integrating planar optode-derived DO distributions with X-ray CT-derived RSA was established. A rootbox equipped with an oxygen-sensitive sensor foil was developed (Figs. 1A, S2A, B). A rice seedling (cv. ‘KSH’) was cultivated in flooded paddy soil under reduced conditions, as indicated by high ferrous iron concentrations in the soil water (Fig. S8). Rhizosphere DO distribution was visualized daily using a planar optode (Figs. 1B, S3A), followed by visualization of RSA in relation to the sensor foil using X-ray CT (Fig. 1C). The optode image was converted into a DO distribution map and overlaid with CT-derived root images (Fig. 1D). The merged images revealed higher DO concentrations (>10%) around shorter roots, whereas DO levels remained relatively low in the background soil and around longer roots.

**Figure 1.**
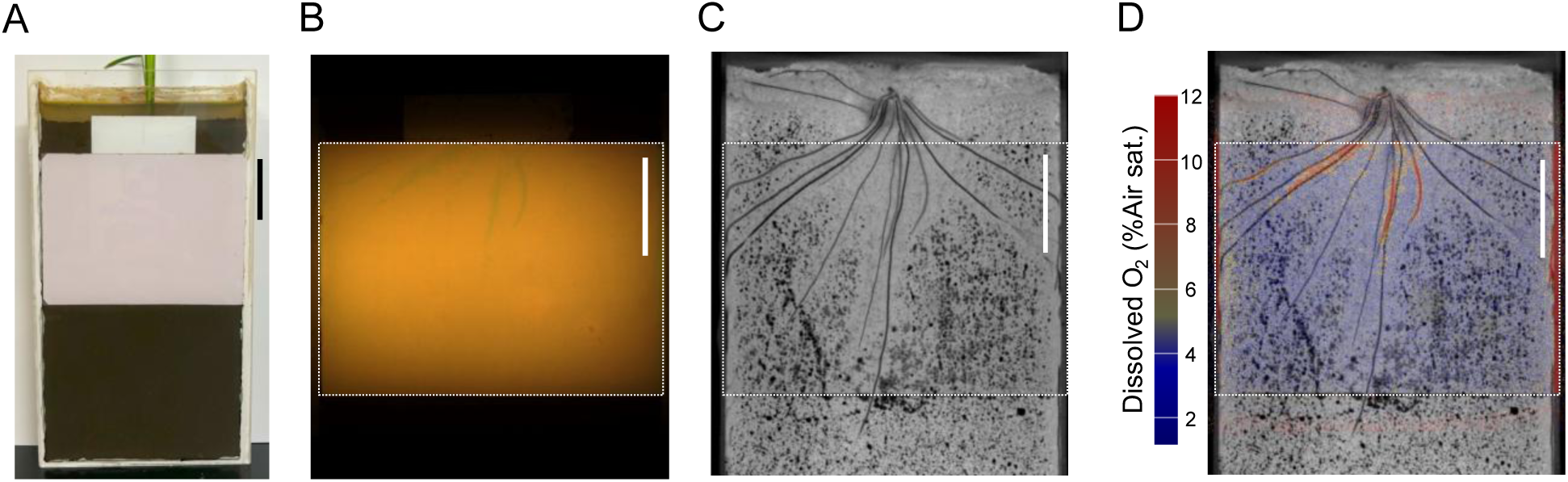
Visualization of rhizosphere oxidation using a rootbox system. **(A)** Front view of the root box. The pink area indicates the oxygen-sensitive optode foil attached to the inner surface of the glass wall. A white laminated tape was placed along the upper edge of the optode foil to prevent root elongation along this boundary. **(B)** RGB image of the optode foil acquired using the camera system. **(C)** Root system image of ‘Koshihikari’ (KSH) at 23 days after sowing visualized by X-ray CT. **(D)** Merged image showing dissolved oxygen (DO) distribution, converted from the optode RGB image in **(B)**, overlaid with the root system image in **(C)**. Colors indicate DO concentrations. White dotted lines indicate the positions of the optode foil in **(B)**–**(D)**. Scale bars = 3 cm.

As part of this workflow, a custom image analysis procedure was developed to quantify DO profiles along crown-root axes (Fig. S4). DO values were extracted along the crown root axes of target roots using a custom software tool (RG2DO-Root) from merged planar optodes and CT images (Fig. S6B). The DO distribution patterns in the representative roots were compared using various circular averaging radii to determine an appropriate spatial averaging scale. Averaging within a 10-pixel radius (∼1.2 mm) produced smoother and less variable DO profiles than smaller radii (Fig. S6A–C). This pattern was consistent with the quantitative noise analysis, indicating that a 10–pixel integration radius reduced the upper noise boundary (95th percentile) relative to point–wise measurements (Fig. S7). Accordingly, DO values averaged within a 10–pixel radius were used in subsequent analyses.

### Developmental changes in rhizosphere oxidation along individual crown roots by daily multimodal imaging

Temporal changes in DO distribution were examined along a representative crown root. At 1 day after emergence (DAE), the crown root measured 42.7 mm in length, with DO values of 5–10% in the mid-root region, specifically between 20 and 30 mm from the base (Fig. 2A, B). This was followed by a sharp decline within approximately 5 mm of the root tip. At 2 DAE, the root elongated by an additional 33.2 mm; DO levels in the region present at 1 DAE decreased to background levels, whereas DO in the newly elongated region (5–20 mm from the root tip) remained at approximately 5%. At 3 DAE, the root tip extended beyond the sensor foil, and DO levels in the regions present at 2 DAE decreased to background levels. Areas with DO levels higher than those of the background indicated ROL, whereas reduced DO levels in the basal regions reflected the formation of an ROL barrier. Similar patterns were observed in the two additional roots (Fig. S9). Integrating the planar optode and X-ray CT imaging enabled the visualization and quantification of daily changes in DO distribution along crown roots.

**Figure 2.**
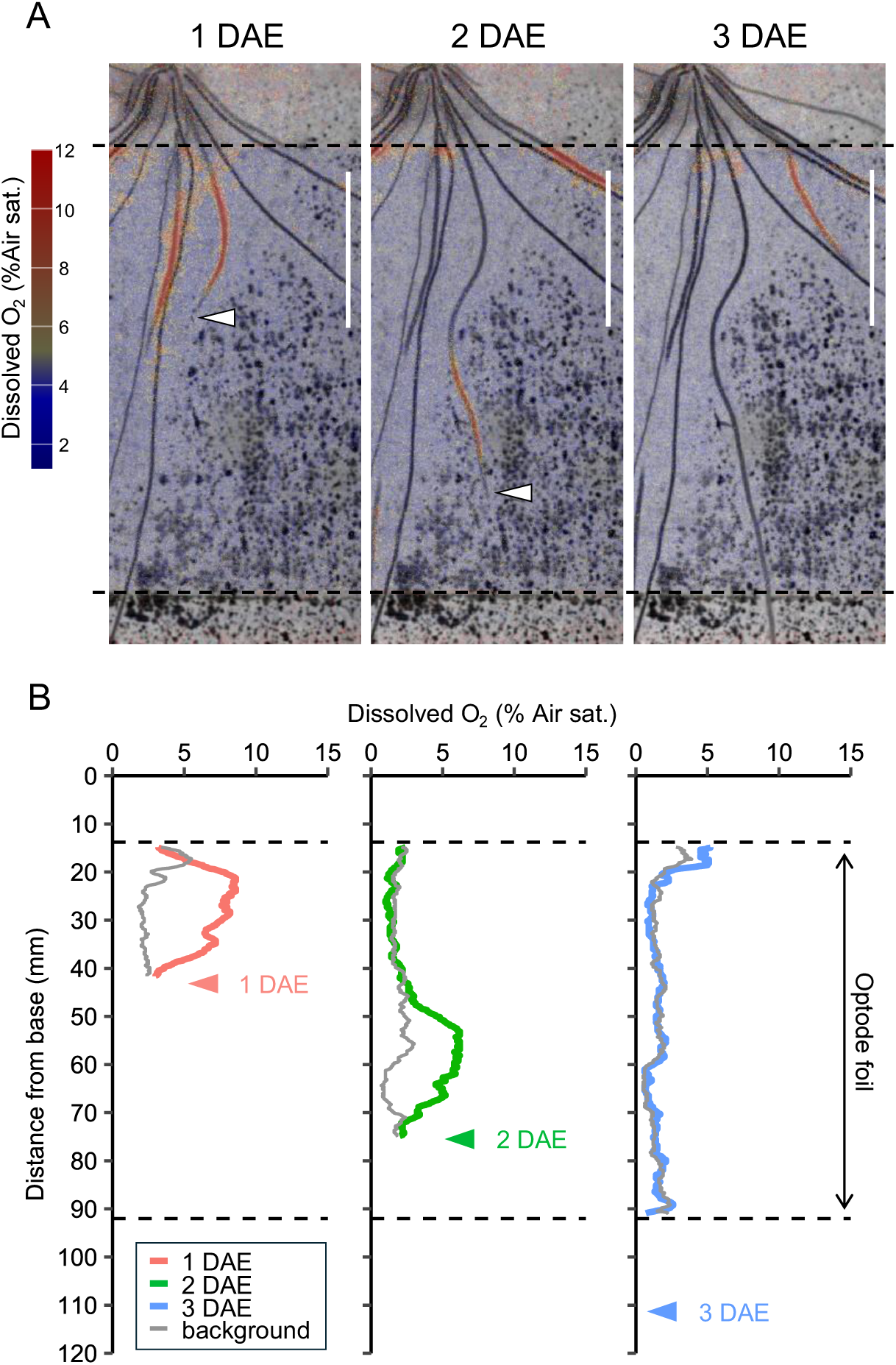
Visualization and quantification of daily changes in longitudinal dissolved oxygen (DO) profiles along a crown root of KSH at 23–25 days after sowing. **(A)** Spatiotemporal visualization of DO concentrations along the crown root. Colors indicate DO concentrations. Scale bars = 3 cm. **(B)** Quantified longitudinal DO profiles corresponding to **(A)**. Colored lines indicate rhizosphere DO values, whereas gray lines represent background DO values. Arrowheads indicate root tip positions. Dashed lines denote the upper and lower boundaries of the optode foil. DAE: days after emergence.

### Distinct root architectural traits among the four rice cultivars

The four rice cultivars (KSH, Muha, QZZ, and Tupa121-3) were compared to assess genotypic variations in root architectural traits associated with rhizosphere oxidation. Shoot and root growth were initially analyzed at an early stage using rhizobox-grown plants. At 21 DAS, the leaf age was comparable among the four cultivars (Fig. S10A); however, the shoot dry weight was lower in KSH and Tupa121-3, with Tupa121-3 also exhibiting a shorter shoot length (Figs. 3A–D, S10B, C). The root dry weight was higher in QZZ than that in KSH, with QZZ exhibiting more crown roots and a longer total root length (Figs. 3E–K, S11A, B). Tupa121-3 produced the fewest crown roots (Fig. 3J). The root cone angle was larger in QZZ and smaller in Muha and Tupa121-3 (Fig. S11D–H). These results demonstrate substantial genotypic variation in root system size and architecture among the four cultivars.

**Figure 3.**
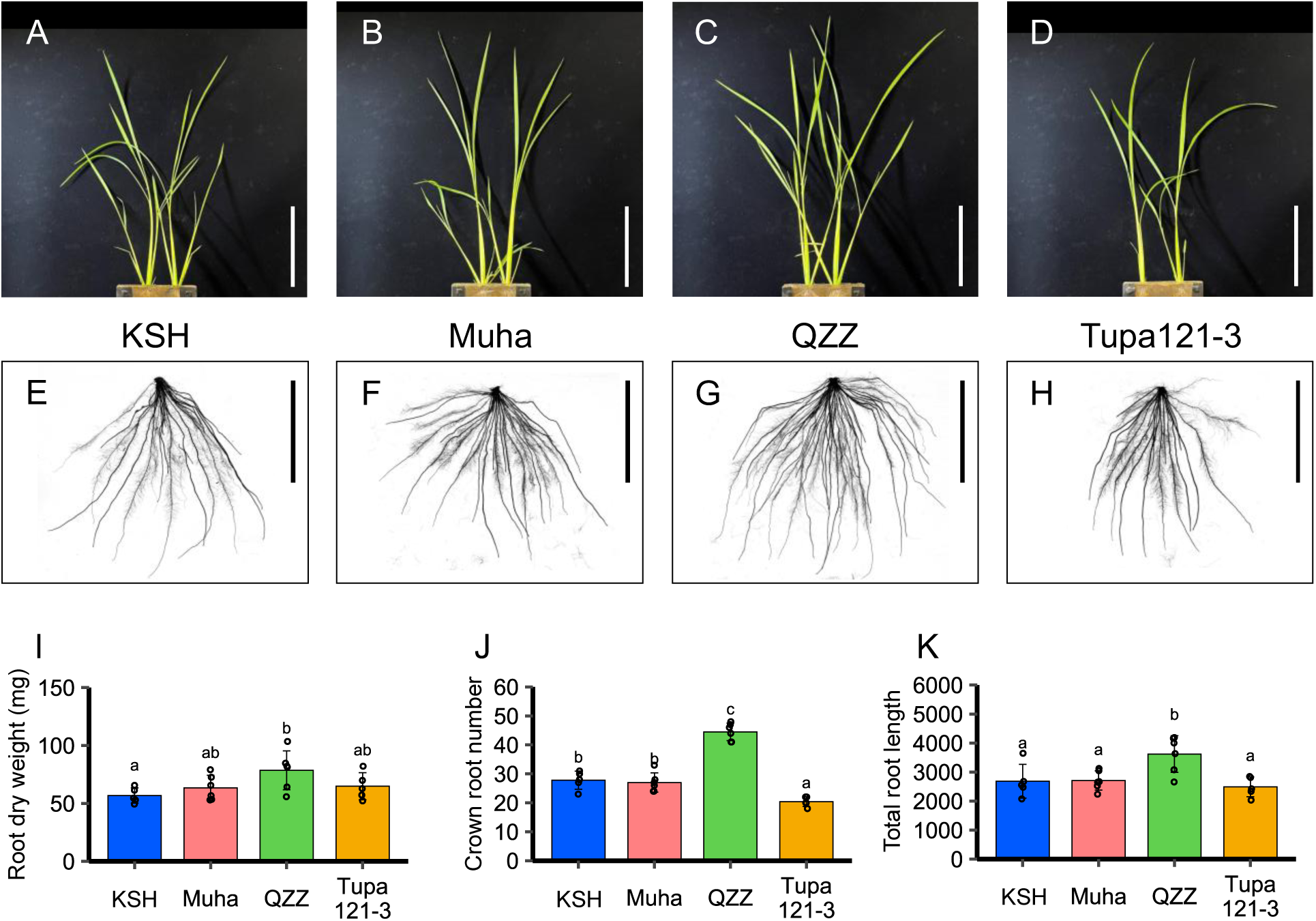
Shoot and root traits in four rice cultivars at 21 days after sowing. **(A–D)** Representative shoot images of each cultivar. Scale bars = 10 cm. **(E–H)** Representative grayscale root system images of each cultivar. Scale bars = 10 cm. **(I–K)** Quantification of shoot and root traits in each cultivar. Bar plots represent mean ± SD (*n* = 5 or 6 plants). Different letters indicate significant differences among cultivars (*P* < 0.05, Tukey’s multiple comparison test).

### Genotypic differences in rhizosphere oxidation patterns associated with root length and age

To characterize the genotypic variation in rhizosphere oxidation patterns, rhizosphere DO distributions were analyzed across the four cultivars as a function of root length and developmental age. Across all cultivars, DO levels near the root tips (0–30 mm) decreased with root elongation, with a particularly pronounced decline observed in shorter roots up to approximately 50 mm in length (Fig. 4). Because DO patterns were closely associated with root growth, as observed in KSH (Fig. 2A, B), roots were classified by length to enable comparative analysis.

**Figure 4.**
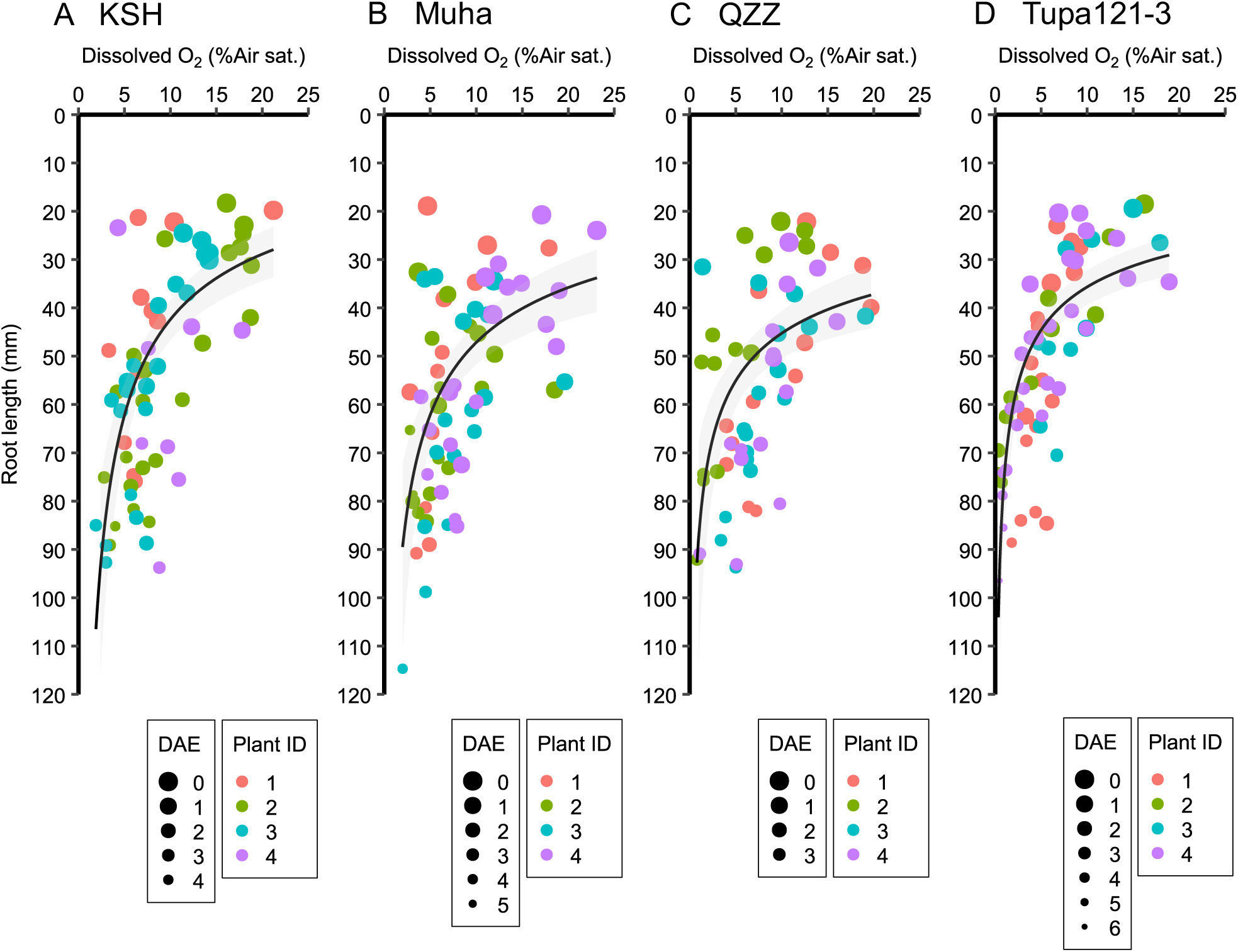
Relationship between crown root length and maximum dissolved oxygen levels in the root tip region (0–30 mm) across four rice cultivars. Each point represents an individual measurement, with colors indicating different plant individuals. Point size denotes root age (days after emergence, DAE). Regression curves were fitted using a log–log linear model. Shaded areas indicate 95% confidence intervals back-transformed from the log scale.

In shorter roots (40–60 mm), rhizosphere DO values were higher than background values at approximately 5–20 mm from the root tip in all four cultivars (Fig. 5E). In the middle-length roots (60–80 mm), rhizosphere DO values were relatively lower than those in shorter roots; however, they remained higher than those of the background, except in Tupa121-3 (Fig. 5F). In Muha, elevated DO levels extended up to 40 mm from the root tips. In longer roots (80–100 mm), elevated rhizosphere DO was confined to the apical regions in KSH and QZZ (up to 20 mm from the root tips), whereas this region was relatively larger in Muha (up to 40 mm) (Fig. 5G). In contrast, no clear increase in rhizosphere DO above the background levels was detected in Tupa121-3. Accordingly, the difference between rhizosphere and background DO was consistently smaller in Tupa121-3 than in KSH across all root-length classes, whereas that in Muha and QZZ was comparable to that in KSH (Fig. S12A–C).

**Figure 5.**
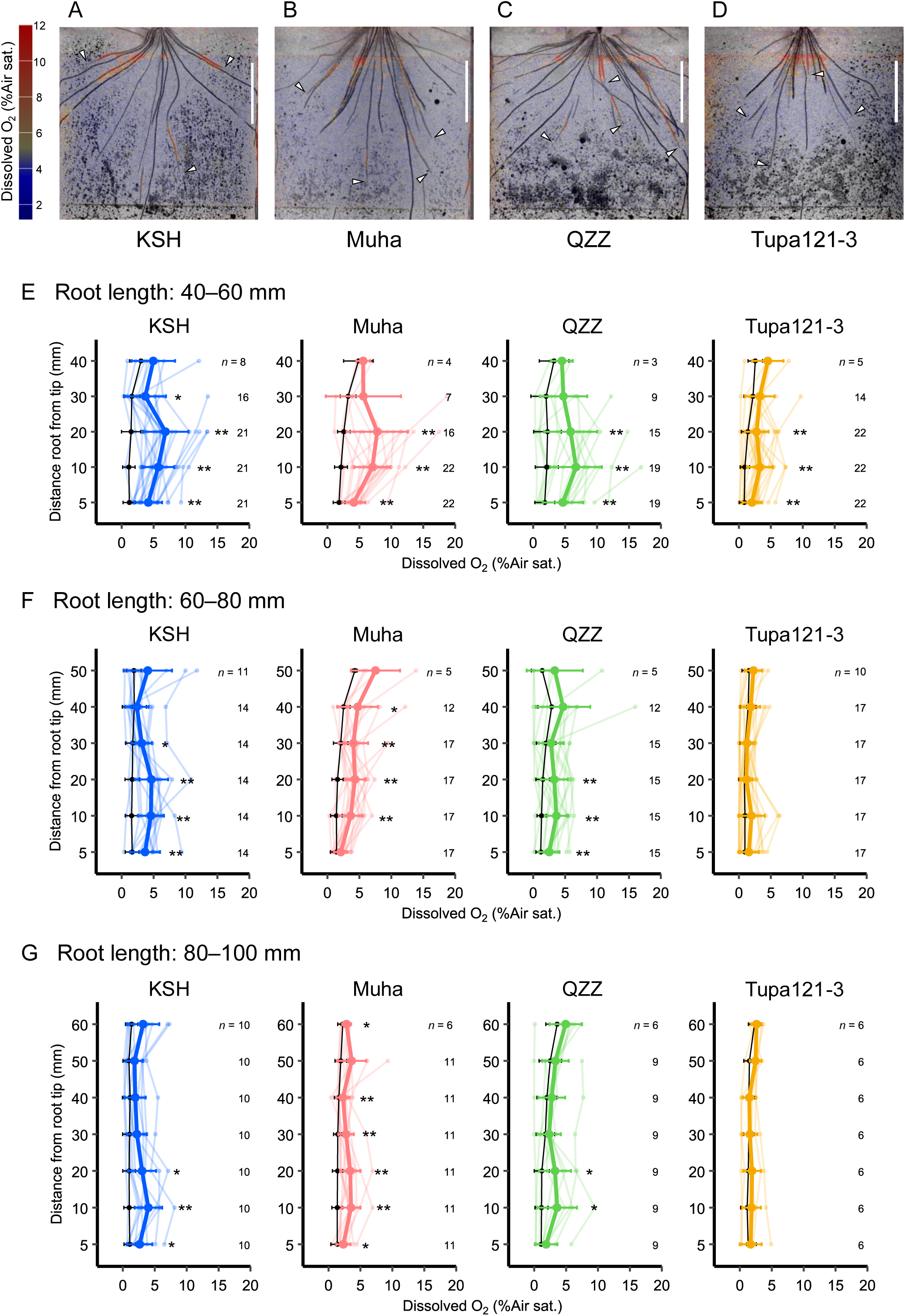
Visualization and quantification of dissolved oxygen (DO) profiles along crown roots in four rice cultivars. **(A–D)** Representative images showing DO distributions in the root systems of each cultivar at 3 weeks after sowing. Arrowheads indicate the positions of elongating root tips. Scale bars = 3 cm. **(E–H)** Longitudinal DO profiles across different root-length classes for each cultivar. Colored lines indicate mean rhizosphere DO values, pale-colored lines indicate values for individual roots, and black lines represent background DO values. Asterisks denote significant differences between rhizosphere and background values (two-tailed Student’s *t* test; *P* < 0.05, *P* < 0.01). The numbers adjacent to the data points indicate the number of replicates.

Subsequently, the impact of root age on the genotypic variations in DO patterns along the root axis was investigated. Root growth rates were lower in QZZ and Tupa121-3 than those in KSH, resulting in developmentally older roots in Tupa121-3 than those in KSH at comparable lengths across the three root-length classes (Fig. S13A, B). Roots in the 60–80 mm length class were subdivided into two age groups (2 and 3 DAE) to more closely examine age-related differences among cultivars independent of root length (Fig. S13C). In KSH, the rhizosphere DO levels tended to be lower in older roots (3 DAE) than those in younger roots (2 DAE). In contrast, the DO levels in Tupa121-3 remained constant between 2 and 3 DAE, suggesting that the influence of root age on DO patterns differed among cultivars. In Tupa121-3, the DO levels were lower than those in KSH in the 10–30 mm region from the root tips at 2 DAE, whereas no clear difference was observed at 3 DAE (Fig. S13C). These results suggest that the reduced rhizosphere oxidation observed in Tupa121-3 can be partially attributed to the differences in developmental timing relative to KSH. However, root age alone did not fully account for the observed patterns, implying that additional cultivar-specific mechanisms regulating ROL could be involved.

### Root thickness was positively associated with rhizosphere oxidation

Given that root diameter affected ROL levels near root tips (Yamauchi et al., 2019), this relationship was analyzed across the four cultivars. Crown root thickness varied across cultivars, with Tupa121-3 displaying the largest root diameter (Figs. 6A–E, S14A). Tupa121-3 possessed a larger stele diameter and cortex width, in addition to a greater number of cortical layers and larger cortical cells than those of the other cultivars (Fig. S14B–F). The aerenchyma area was larger in Tupa121-3 at various positions along the root axis, whereas the proportion of aerenchyma relative to the cortex was comparable to that of the other cultivars (Fig. S14G–I).

**Figure 6.**
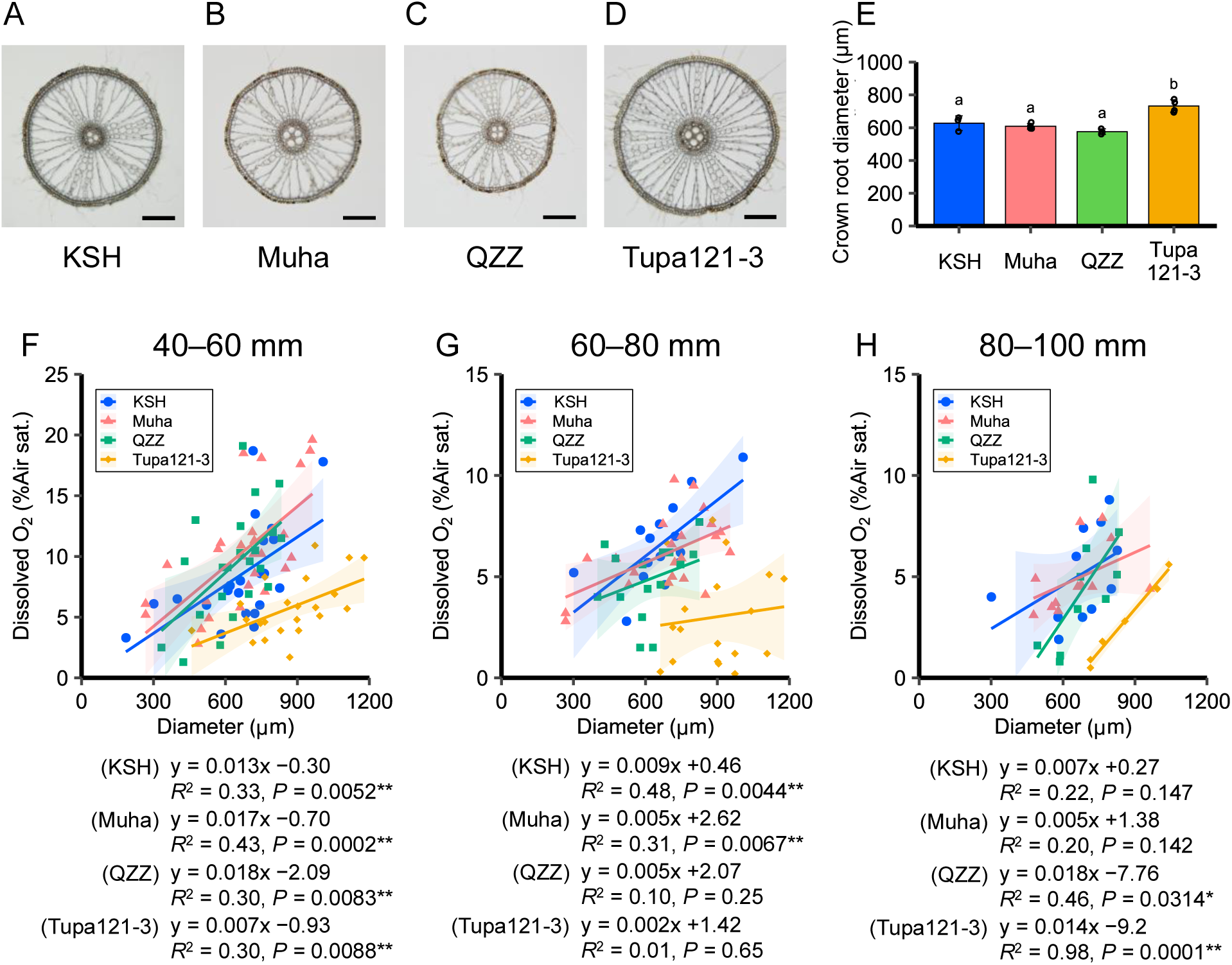
Effects of crown root diameter on dissolved oxygen (DO) levels near the root tip. **(A–D)** Representative cross-sectional images of crown roots at 50 mm from the root tip for each cultivar. Scale bars = 200 µm. **(E)** Average crown root diameter of emerged roots at 21 days after sowing. Bar plots represent mean ± SD (*n* = 3 or 4 plants). Different letters indicate significant differences among cultivars (*P* < 0.05, Tukey’s multiple comparison test). **(F–H)** Relationships between crown root diameter and maximum DO levels in the root tip region (0–30 mm) for each cultivar across different root-length classes. The numbers above the panels indicate the corresponding root-length classes. Points represent individual measurements, with colors indicating different cultivars. Solid lines indicate cultivar-specific fitted regression lines estimated using linear models (DO ∼ crown root diameter), and shaded areas denote 95% confidence intervals. Regression slopes and coefficients of determination (*R*^2^) were calculated for each cultivar (Pearson’s correlation analysis, *P* < 0.05, *P* < 0.01).

The relationship between root thickness and DO levels near the root tips was analyzed to assess the influence of the observed differences in crown root thickness. The crown root diameter was positively correlated with the DO levels near the root tips across all cultivars, with a notably strong relationship observed in the short-root class (Fig. 6F–H). Although Tupa121-3 followed the same positive trend as the other cultivars, it consistently demonstrated lower DO values despite its thicker roots. These results indicate that increased root diameter is associated with higher DO levels near the root tip. Additional factors are likely to account for the low DO observed in Tupa121-3.

## Discussion

### Multimodal imaging bridges solution-based ROL studies to dynamic rhizosphere oxidation patterns in soil

In this study, we developed a multimodal visualization system integrating planar optodes and X-ray CT to investigate rhizosphere oxidation in rice under reduced soil conditions. By developing a custom analysis pipeline to quantify oxygen profiles along individual root axes, this system enabled non-destructive daily time-course tracking of rhizosphere oxygen dynamics in relation to root elongation, aging, and thickness. Analyses using the multimodal visualization system revealed that ROL patterns in soil exhibited dynamic changes in response to root elongation. Across the tested cultivars, younger and shorter roots (40–60 mm) demonstrated relatively higher DO levels over large root areas, whereas in longer roots (60–80 and 80–100 mm), elevated DO was spatially restricted toward the root tip (Figs. 2, 5E–G, and S9). This pattern is consistent with that of previous findings from stagnant solution experiments, indicating that ROL barrier formation occurs in more mature root regions, thereby restricting oxygen leakage to the apical zones (Colmer, 2006; Ejiri et al., 2021).

Although longitudinal ROL patterns have been extensively studied, less attention has been given to how ROL intensity at the root tip changes during individual root elongation. Across all cultivars tested in this study, shorter roots exhibited a steeper rate of decline in DO at the root tip, indicating that early root elongation represents a particularly dynamic phase of rhizosphere oxygen regulation (Fig. 4). Therefore, integrating X-ray CT-based root tracking with planar optode imaging provides a powerful approach for capturing the developmental dynamics of rhizosphere oxidation in soil.

### Genotypic differences in the magnitude of rhizosphere oxidation associated with root elongation and developmental age

We observed genotypic variations in ROL patterns among rice cultivars. Previous studies utilizing stagnant solution systems have demonstrated genotypic differences in ROL barrier formation based on longitudinal ROL distributions (Colmer et al., 1998; Colmer, 2003b). Here, ROL was inferred from differences between rhizosphere and background DO levels near root tips (Figs. 5E–G, S12). The absence of detectable increases in rhizosphere DO in basal regions for at least one root-length class across all tested cultivars (Fig. 5E–G) suggests the formation of an ROL barrier that restricts oxygen leakage from mature root tissues. In contrast, the elevation of rhizosphere DO near the root tips was consistently lower in Tupa121-3 than that in KSH across all root-length classes (Fig. S12A–C). Moreover, no statistically significant increase in DO was observed at the root tips of Tupa121-3 in the longer roots (60–80 and 80-100 mm; Fig. 5F, G). Notably, the longer roots maintained comparable root elongation rates (Fig. S13B), indicating that sufficient oxygen was supplied internally to support root growth, despite the low rhizosphere oxygen signal. Tupa121-3 demonstrated lower root elongation rates than those in KSH, leading to relatively older root ages within the same root-length classes (Fig. S13A, B). In KSH, older roots within the same root-length class tended to show lower DO levels than younger roots, with the DO levels of these older KSH roots being comparable to those observed in Tupa121-3 (Fig. S13C). However, in younger roots, the DO levels in Tupa121-3 were significantly lower than those in KSH, indicating that differences in root age alone do not fully account for the reduced DO levels in Tupa121-3. These results suggest that, in addition to developmental timing, other genotype–specific physiological or anatomical factors contribute to the observed ROL patterns.

### Root thickness contributes to rhizosphere oxidation but is insufficient to explain genotypic variation

Within a genotype, thicker roots tended to exhibit higher rhizosphere DO levels near root tips (Fig. 6F–H), likely reflecting improved internal oxygen transport capacity. Rice plants typically produce both thin and thick roots depending on nodal origin (Kawata et al., 1963). Yamauchi et al. (2019) used a stagnant solution system and found that although the longitudinal patterns of ROL were similar between thin and thick roots, the magnitude of ROL near the root tips was higher in thick roots. They attributed this difference to anatomical traits, including a higher aerenchyma-to-cortex ratio near the root tips and larger cortex and aerenchyma areas in thicker roots, thereby indicating greater internal oxygen transport capacity. Among the genotypes examined in this study, Tupa121-3 exhibited thicker crown roots than those of the other genotypes, along with increased cortical and aerenchymal areas (Figs. 6A–E, S14). However, despite these anatomical features, the DO levels near the root tips were relatively lower in Tupa121-3 (Figs. 6F–H, S12). This contrast indicates that among genotypes, root thickness alone does not directly translate into greater oxygen leakage to the rhizosphere. This discrepancy likely reflects the influence of additional anatomical and physiological traits, along with rhizosphere conditions, including microbial communities, that differ between stagnant solutions and soil environments.

### Mechanistic implications for genotypic differences in rhizosphere oxidation

To understand the basis for the reduced rhizosphere DO levels in Tupa121–3 despite its thicker roots, we next considered differences in internal root anatomy and oxygen consumption. In Tupa121-3, the stele diameter was larger than that in the other genotypes (Fig. S14C). In *Urochloa humidicola*, cellular-scale partial pressure of oxygen (pO_2_) measurements using Clark-type O_2_ microsensors and a root-sleeving electrode have revealed higher oxygen consumption and greater liquid-phase diffusive resistances in stele and exodermal/epidermis cells than in gas-filled aerenchyma (Jiménez et al., 2024b). In this study, cross-sectional analyses along the longitudinal root axis indicated that suberin lamella formation was absent at 10 and 20 mm from the root tips (Fig. S15), implying relatively high oxygen permeability in these apical regions. Under these conditions, increased respiratory activity in the stele and exodermal/epidermal cells may enhance internal oxygen consumption, thereby decreasing oxygen leakage to the rhizosphere despite sufficient internal oxygen supply levels to sustain root elongation. Alongside root anatomical and physiological factors, microbial processes may contribute to the observed genotypic variations in rhizosphere DO distribution. Aerobic microorganisms, including methanotrophs, nitrifiers, and iron-oxidizing bacteria, inhabit the rice rhizosphere (Ding et al., 2019). Genotype-dependent variations in microbial community composition and activity can influence local oxygen dynamics. Further studies integrating measurements of root respiration, internal oxygen distribution, and rhizosphere microbial community analyses are necessary to identify the mechanisms underlying genotypic differences in ROL more precisely.

### Technical advantages and limitations of the multimodal imaging system

The multimodal system developed here offers a high-throughput framework compared with single-point measurements, requiring less than 1 min per rootbox for optode imaging and 10 min for X-ray CT scanning of four rootboxes. This efficiency enabled the simultaneous analysis of multiple genotypes to elucidate genotypic spatiotemporal differences in rhizosphere oxygen distribution. As the system combines commercially available imaging components, it can be readily adopted by other research groups. Notably, the planar optode setup allowed visualization across a large field of view (100 mm width × 75 mm depth), enabling the tracking of individual crown roots exceeding 80 mm in length. Simultaneously, the spatial noise inherent in the optode foil necessitated normalization by spatial averaging, resulting in an effective spatial resolution of approximately 2 mm in diameter when using circular regions with a radius of 10 pixels (Figs. S6, S7). This integrated approach provides a practical and scalable framework for linking root development to the rhizosphere oxidation dynamics in flooded soils.

Despite its advantages, our system had several limitations that warrant further consideration. In this study, the crown root axes were examined to analyze spatiotemporal changes in rhizosphere oxidation associated with individual root development. Zones with elevated oxygen concentrations have been reported to form near the base of plants, where high densities of small, young crown roots and lateral roots are present (Larsen et al., 2015). Rice produces two types of lateral roots characterized by differing anatomical structures, including the presence or absence of aerenchyma and ROL barrier formation, leading to distinct ROL patterns (Kawai et al., 2017, 2022; Noorrohmah et al., 2020). Higher-resolution X-ray CT scanning is necessary to accurately visualize lateral root development and spatial distribution. This scanning requires longer acquisition times and a narrower region of interest, which conflicts with the high-throughput characteristics of the present framework. Nevertheless, lateral roots constitute a substantial proportion of the rice root system, and their cumulative contribution may influence overall plant rhizosphere oxidation.

### Implications and future perspectives for rhizosphere oxygen dynamics

Given the characteristics of the developed multimodal system discussed above, integrating multi-scale quantification of DO distributions may enhance understanding of rhizosphere oxidation processes. Previous studies have demonstrated that ROL from the total rice root system, quantified using the Ti^3+^-citrate method, exhibits a negative correlation with methane emissions under reduced soil conditions (Zheng et al., 2018; Chen et al., 2019; Qi et al., 2024). The oxygen released from rice roots can suppress methane production by inhibiting anaerobic methanogenesis and may promote methane oxidation in the oxic microsites surrounding the roots (Van Bodegom et al., 2001; Li et al., 2022). In these studies, ROL was generally positively correlated with the total root volume; however, the contribution of the spatial distribution of oxygen release within the root system has not been quantified. Further integration of the developed multimodal system with approaches that quantify RSA, total ROL, and process-based modeling frameworks may be required to comprehensively extend these findings to whole-plant oxygen release, soil oxidation, and ecosystem-scale processes. This multi–scale integration may enhance our understanding of the collective influence of root developmental traits on rhizosphere oxygen environments under field-relevant conditions.

## Conclusions

This study revealed the spatiotemporal dynamics of rhizosphere oxygen associated with individual root growth and developmental traits using a multimodal imaging approach. Previous studies using planar optodes to visualize rhizosphere oxygen distribution have primarily focused on the spatial extent of root-induced oxic zones and their responses to shoot environmental conditions (Larsen et al., 2015; Lenzewski et al., 2018; Koop-Jakobsen et al., 2018; Han et al., 2019; Maisch et al., 2020). In contrast, spatiotemporal changes in the ROL profiles during root development have primarily been inferred from single-point measurements conducted in hydroponic or stagnant solution systems in rice (e.g., Colmer, 2003b). These studies have demonstrated that ROL patterns are influenced by the root developmental stage, oxygen availability, and phytochemical conditions in the surrounding solution, thereby reflecting associated histochemical changes in root tissues (Colmer et al., 2019; Jiménez et al., 2025). In this context, rhizosphere oxidation has been hypothesized to be closely associated with root growth and morphological or anatomical characteristics. By integrating planar optodes with X-ray CT, rhizosphere oxygen dynamics were directly visualized under reduced soil conditions, revealing how they change in concert with individual root growth and developmental traits. This multimodal approach bridges the gap between solution–based inferences and soil–based observations, thereby providing a framework for linking root developmental traits with spatiotemporal patterns of rhizosphere oxidation under field–relevant conditions.

## Supporting information

RG2DO-Root

## Acknowledgments

We thank H. Imaizumi (NIAS, NARO) and N. Tanaka (NICS, NARO) for their technical support with the microscopy experiments; Y. Fukuda, H. Tanaka, K. Onoe, N. Kanno, and C. Kawashima for their technical assistance with plant phenotyping; K. Kambe and A. Sugano for the field observation; M. Tsunematsu, Y. Kajihara for the soil chemical analysis; K. Shiono (Fukui Prefectural U.) and M. Ejiri (GRA&GREEN Inc.) for their valuable advice on the planar optode setup and insightful discussions. Fe (II) concentration measurements were performed at the Advanced Analysis Center of NARO.

## Author Contributions

T.K., T.T., and Y.U designed the study. T.K., X.M, D.F., K.K.H., and S.M.K. conducted the experiments. T.K., S.T., X.M., and D.F. analyzed the data. T.K. and Y.U. drafted the original manuscript, and all authors commented on and approved the final version for submission.

## Supplementary data

The following materials are available in the online version of this article:

**Figure S1.** Root systems of four rice cultivars cultivated in a paddy field at different developmental stages.

**Figure S2.** Rootbox system, cultivation conditions, and imaging platforms used in this study.

**Figure S3.** Cultivation conditions in this study.

**Figure S4.** Workflow for multimodal data collection and analysis of rhizosphere dissolved oxygen (DO) distribution and root development.

**Figure S5.** Calibration of oxygen-sensitive optode foils before and after measurements. **Figure S6.** Comparison of dissolved oxygen (DO) profiles along a crown root obtained using different circular averaging radii in ‘Koshihikari’ (KSH) at 24 days after sowing.

**Figure S7.** Sensor noise characteristics of planar optodes and noise reduction by spatial averaging.

**Figure S8.** Ferrous iron concentrations in the soil pore water of a blank rootbox (without plants) and rootboxes planted with different cultivars at 20 days after sowing.

**Figure S9.** Daily changes in longitudinal dissolved oxygen (DO) profiles along two crown roots of KSH at 18–21 days after sowing.

**Figure S10.** Shoot traits of four rice cultivars at 21 days after sowing.

**Figure S11.** Root traits of four rice cultivars at 21 days after sowing.

**Figure S12.** Longitudinal profiles of differences in dissolved oxygen (DO) levels between the rhizosphere and background along crown roots across different root-length classes for four rice cultivars.

**Figure S13.** Effects of root age on dissolved oxygen (DO) profiles in four rice cultivars.

**Figure S14.** Anatomical structure of crown roots in four rice cultivars at 21 days after sowing.

**Figure S15.** Suberin lamella formation at different distances from the root tip in a crown root of ‘Tupa121-3’.

## Data availability

The source code is available in Supplemental Data RG2DO-Root.zip.

## References

Armstrong W, Wright EJ. 1975. Radial oxygen loss from roots: The theoretical basis for the manipulation of flux data obtained by the cylindrical platinum electrode technique. Physiol Plant. 35:21–26. 10.1111/j.1399-3054.1975.tb03861.x.

Armstrong W. 2000. Oxygen distribution in wetland plant roots and permeability barriers to gas-exchange with the rhizosphere: a microelectrode and modelling study with *Phragmites australis*. Ann Bot. 86:687–703. 10.1006/anbo.2000.1236.

Blossfeld S. 2013. Light for the dark side of plant life: —Planar optodes visualizing rhizosphere processes. Plant Soil. 369:29–32. 10.1007/s11104-013-1767-0.

Chen Y et al. 2019. Rice root morphological and physiological traits interaction with rhizosphere soil and its effect on methane emissions in paddy fields. Soil Biol Biochem. 129:191–200. 10.1016/j.soilbio.2018.11.015.

Colmer TD, Gibberd MR, Wiengweera A, Tinh TK. 1998. The barrier to radial oxygen loss from roots of rice (*Oryza sativa* L.) is induced by growth in stagnant solution. J Exp Bot. 49:1431–1436. 10.1093/jxb/49.325.1431.

Colmer TD. 2003a. Long–distance transport of gases in plants: a perspective on internal aeration and radial oxygen loss from roots. Plant Cell Environ. 26:17–36. 10.1046/j.1365-3040.2003.00846.x.

Colmer TD. 2003b. Aerenchyma and an inducible barrier to radial oxygen loss facilitate root aeration in upland, paddy and deep-water rice (*Oryza sativa* L.). Ann Bot. 91:301–309. 10.1093/aob/mcf114.

Colmer TD, Cox MCH, Voesenek LACJ. 2006. Root aeration in rice (*Oryza sativa*): evaluation of oxygen, carbon dioxide, and ethylene as possible regulators of root acclimatizations. New Phytol. 170:767–778. 10.1111/j.1469-8137.2006.01725.x.

Colmer TD, Pedersen O. 2008. Oxygen dynamics in submerged rice (*Oryza sativa*). New Phytol. 178:326–334. 10.1111/j.1469-8137.2007.02364.x.

Colmer TD et al. 2019. Rice acclimation to soil flooding: Low concentrations of organic acids can trigger a barrier to radial oxygen loss in roots. Plant Cell Environ. 42:2183–2197. 10.1111/pce.13562.

Ding LJ et al. 2019. Microbiomes inhabiting rice roots and rhizosphere. FEMS Microbiol Ecol. 10.1093/femsec/fiz040

Drew MC, Lynch JM. 1980. Soil anaerobiosis, microorganisms, and root function. Annu Rev Phytopathol. 18:37–66. 10.1146/annurev.py.18.090180.000345.

Ejiri M, Fukao T, Miyashita T, Shiono K. 2021. A barrier to radial oxygen loss helps the root system cope with waterlogging-induced hypoxia. Breed Sci. 71:40–50. 10.1270/jsbbs.20110.

Fang W et al. 2021. Combining multiple high-resolution *in situ* techniques to understand phosphorous availability around rice roots. Environ Sci Technol. 55:13082–13092. 10.1021/acs.est.1c05358.

Hamamoto S et al. 2025. Seasonal changes in methane emissions via different pathways from a rice paddy field. Paddy Water Environ. 23:333–342. 10.1007/s10333-025-01018-7.

Han C et al. 2019. High-resolution imaging of rhizosphere oxygen (O_2_) dynamics in *Potamogeton crispus*: effects of light, temperature and O_2_ content in overlying water. Plant Soil. 441:613–627. 10.1007/s11104-019-04150-6.

Heeraman DA, Hopmans JW, Clausnitzer V. 1997. Three dimensional imaging of plant roots in situ with X-ray computed tomography. Plant Soil. 189:167–179. 10.1023/B:PLSO.0000009694.64377.6f.

Hothorn T, Bretz F, Westfall P. 2008. Simultaneous inference in general parametric models. Biometrical J. 50:346–363. 10.1002/bimj.200810425.

Iqbal MF et al. 2021. Limited aerenchyma reduces oxygen diffusion and methane emission in paddy. J Environ Manage. 279:111583. 10.1016/j.jenvman.2020.111583.

Jiménez J de la C, Pellegrini E, Pedersen O, Nakazono M. 2021. Radial oxygen loss from plant roots—Methods. Plants. 10:2322. 10.3390/plants10112322.

Jiménez J de la C et al. 2024a. *Leaf Gas Film 1* promotes glycerol ester accumulation and formation of a tight root barrier to radial O_2_ loss in rice. Plant Physiol. 196:2437–2449. 10.1093/plphys/kiae458.

Jiménez J de la C, Armstrong W, Colmer TD, Pedersen O. 2024b. Overcoming constraints to measuring O_2_ diffusivity and consumption of intact roots. Plant Physiol. 195:283–286. 10.1093/plphys/kiae046.

Jiménez J de la C et al. 2025. Formation of apoplastic barriers to radial O_2_ loss in rice roots: effects of low-O_2_ and high–Fe conditions, and the roles of suberin, glycerol esters, and iron plaques. Plant Cell Environ. 48:2937–2949. 10.1111/pce.15319.

Kajiura M, Saito M, Ma X, Nishikawa J, Tokida T. 2025. Pathway-specific flux and dissolved CH_4_ pool in the soil across 22 rice cultivars. J Agric Meteorol. 81:57–65. 10.2480/agrmet.D-24-00030.

Kawai T, Nosaka-Takahashi M, Yamauchi A, Inukai Y. 2017. Compensatory growth of lateral roots responding to excision of seminal root tip in rice. Plant Root. 11:48–57. 10.3117/plantroot.11.48.

Kawai T et al. 2022. *WUSCHEL-related homeobox* family genes in rice control lateral root primordium size. Proc Natl Acad Sci U S A. 119:e2101846119. doi: 10.1073/pnas.2101846119.

Kawata S, Yamazaki K, Ishihara K, Shibayama H, Lai K. 1963. Studies on root system formation in rice plants in a paddy. Jpn J Crop Sci. 32:163–180. 10.1626/jcs.32.163.

Koop-Jakobsen K, Mueller P, Meier R, Liebsch G, Jensen K. 2018. Plant-sediment interactions in salt marshes – An optode imaging study of O_2_, pH, and CO_2_ gradients in the rhizosphere. Front Plant Sci. 9:541. 10.3389/fpls.2018.00541.

Kotula L, Ranathunge K, Schreiber L, Steudle E. 2009. Functional and chemical comparison of apoplastic barriers to radial oxygen loss in roots of rice (*Oryza sativa* L.) grown in aerated or deoxygenated solution. J Exp Bot. 60:2155–2167. 10.1093/jxb/erp089.

Larsen M, Borisov S, Grunwald B, Klimant I, Glud R. 2011. A simple and inexpensive high resolution color ratiometric planar optode imaging approach: application to oxygen and pH sensing. Limnol Oceanogr Methods. 9:348–360. 10.4319/lom.2011.9.348.

Larsen M, Santner J, Oburger E, Wenzel W, Glud R. 2015. O_2_ dynamics in the rhizosphere of young rice plants (*Oryza sativa* L.) as studied by planar optodes. Plant Soil. 390:279–292. 10.1007/s11104-015-2382-z.

Lenzewski N et al. 2018. Dynamics of oxygen and carbon dioxide in rhizospheres of *Lobelia dortmanna* – a planar optode study of belowground gas exchange between plants and sediment. New Phytol. 218:131–141. 10.1111/nph.14973.

Li S et al. 2022. Reducing methane emission by promoting its oxidation in rhizosphere through nitrogen-induced root growth in paddy fields. Plant Soil. 474:541–560. 10.1007/s11104-022-05360-1.

Maisch M, Lueder U, Kappler A, Schmidt C. 2019. Iron lung: How rice roots induce iron redox changes in the rhizosphere and create niches for microaerophilic Fe(II)-oxidizing bacteria. Environ Sci Technol Lett. 6:600–605. 10.1021/acs.estlett.9b00403.

Maisch M, Lueder U, Kappler A, Schmidt C. 2020. From plant to paddy—How rice root iron plaque can affect the paddy field iron cycling. Soil Syst. 4:28. 10.3390/soilsystems4020028.

Noorrohmah S, Takahashi H, Nakazono M. 2020. Formation of a barrier to radial oxygen loss in L-type lateral roots of rice. Plant Root. 14:33–41. 10.3117/plantroot.14.33.

Pedersen O, Rich SM, Colmer TD. 2009. Surviving floods: leaf gas films improve O_2_ and CO_2_ exchange, root aeration, and growth of completely submerged rice. Plant J. 58:147–156. 10.1111/j.1365-313X.2008.03769.x.

Qi Z et al. 2024. Effect and mechanism of root characteristics of different rice varieties on methane emissions. Agronomy. 14:595. 10.3390/agronomy14030595.

R Core Team. 2022. R: a language and environment for statistical computing. Vienna, Austria: R Foundation for Statistical Computing. https://www.r-project.org.

Shiono K et al. 2022. Imaging the snorkel effect during submerged germination in rice: Oxygen supply via the coleoptile triggers seminal root emergence underwater. Front Plant Sci. 13:946776. 10.3389/fpls.2022.946776.

Shiono K, Nakazono M. 2024. Development and regulation of a radial oxygen loss barrier to acclimate to anaerobic conditions. responses of plants to soil flooding. In: Sakagami JI, Nakazono M editors. Responses of Plants to Soil Flooding. Springer, p. 119–138. 10.1007/978-981-99-9112-9_8.

Teramoto S et al. 2020. High-throughput three-dimensional visualization of root system architecture of rice using X-ray computed tomography. Plant Methods. 16:66. 10.1186/s13007-020-00612-6.

Uga Y et al. 2009. Variation in root morphology and anatomy among accessions of cultivated rice (*Oryza sativa* L.) with different genetic backgrounds. Breed Sci. 59:87–93. 10.1270/jsbbs.59.87.

Uga Y et al. 2013. Control of root system architecture by *DEEPER ROOTING 1* increases rice yield under drought conditions. Nat Genet. 45:1097–1102. 10.1038/ng.2725.

Van Bodegom P, Goudriaan J, Leffelaar P. 2001. A mechanistic model on methane oxidation in a rice rhizosphere. Biogeochemistry. 55:145–177. 10.1023/A:1010640515283.

Wei T, Simko V. 2021. corrplot: visualization of a correlation matrix. R package version 0.92. https://cran.r-project.org/package=corrplot.

Wei L et al. 2025. Carbon stabilization by iron plaque on rice roots: The role of oxygen loss. Soil Biol Biochem. 210:109947. 10.1016/j.soilbio.2025.109947.

Williams PN et al. 2014. Localized flux maxima of arsenic, lead, and iron around root apices in flooded lowland rice. Environ Sci Technol. 48:8498–8506. 10.1021/es501127k.

Yamauchi T, Colmer TD, Pedersen O, Nakazono M. 2018. Regulation of root traits for internal aeration and tolerance to soil waterlogging-flooding stress. Plant Physiol. 176:1118–1130. 10.1104/pp.17.01157.

Yamauchi T, Abe F, Tsutsumi N, Nakazono M. 2019. Root cortex provides a venue for gas-space formation and is essential for plant adaptation to waterlogging. Front Plant Sci. 10:259. 10.3389/fpls.2019.00259.

Yamauchi T, Li J, Sumi K. 2025. A rapid staining method for the detection of suberin lamellae in the root endodermis and exodermis. Plant Biotechnol. 42:185–188. 10.5511/plantbiotechnology.25.0312a.

Yin Y et al. 2021. Noninvasive imaging of hollow structures and gas movement revealed the gas partial–pressure–gradient–driven long–distance gas movement in the aerenchyma along the leaf blade to submerged organs in rice. New Phytol. 232:1974–1984. 10.1111/nph.17726.

Yoshida S, Forno DA, Cock JH, Gomez KA. 1976. Routine procedure for growing rice plants in culture solution. In: Yoshida S, Forno DA, Cock JH, Gomez KA editors. Laboratory manual for physiological studies of rice. International Rice Research Institute. p. 61–66.

Zheng H, Fu Z, Zhong J, Long W. 2018. Low methane emission in rice cultivars with high radial oxygen loss. Plant Soil. 431:119–128. 10.1007/s11104-018-3747-x.

