## Supplementary_data for "Characterization of Rhizosphere Oxidation Associated with Root Development in Rice Using Planar Oxygen Optodes"

**This PDF file includes:**

Figures S1 to S15

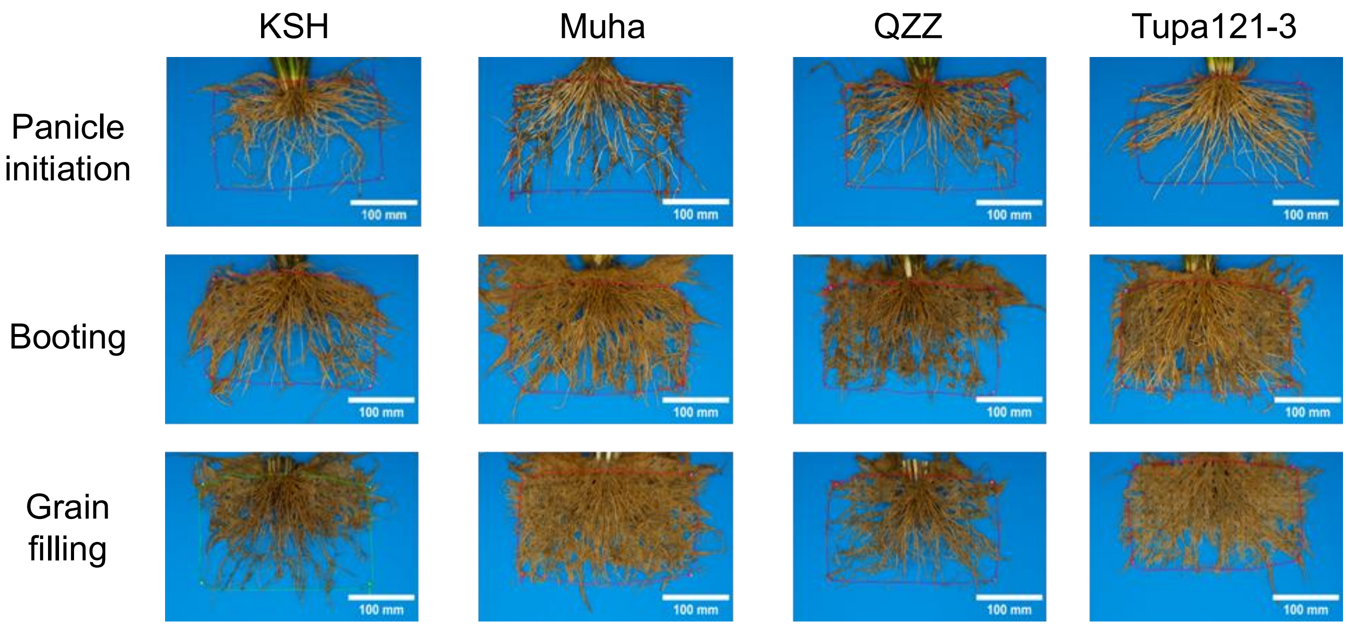


**Figure S1.** Root systems of four rice cultivars cultivated in a paddy field at different developmental stages. Representative images of root systems are shown for each cultivar.

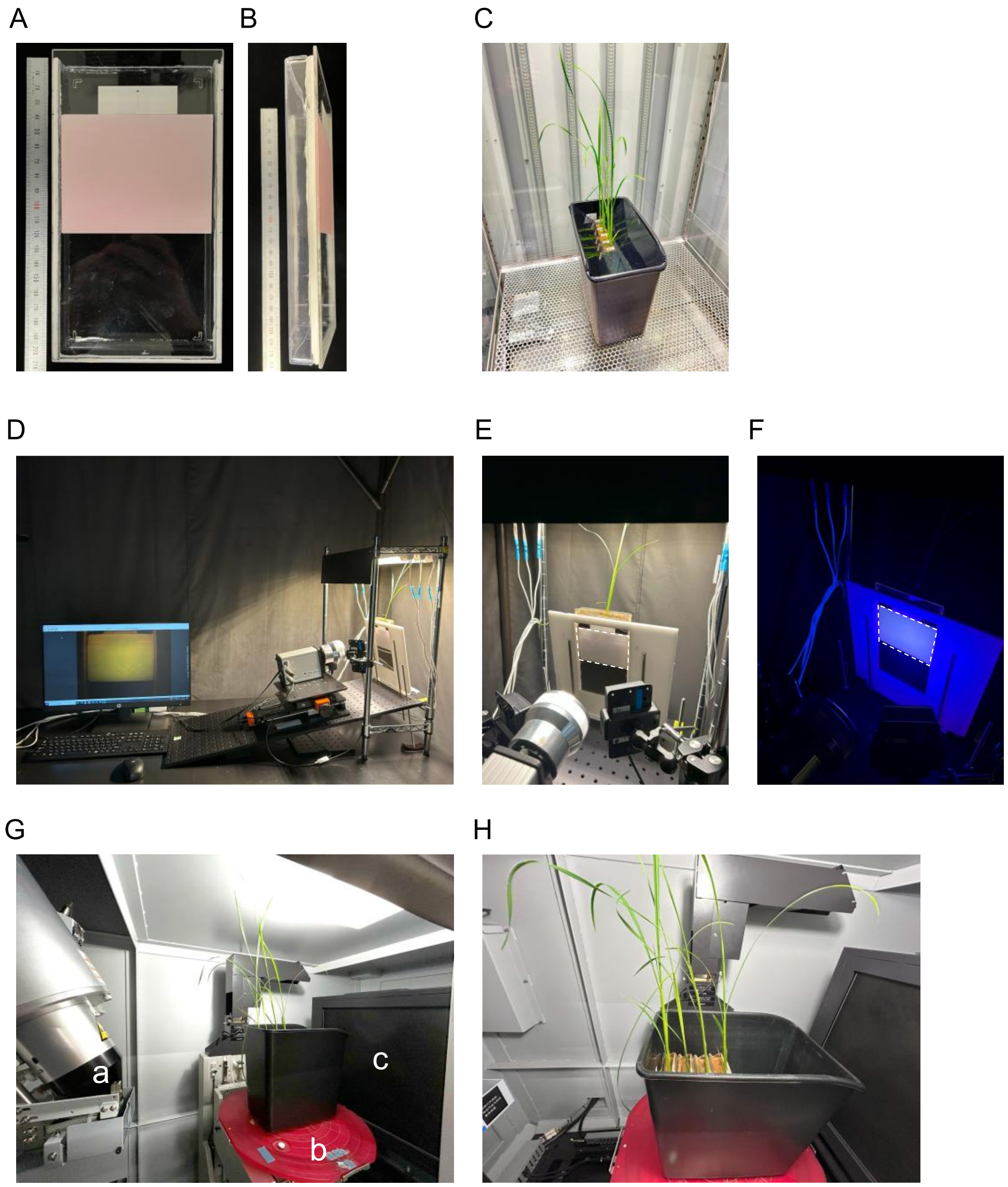


**Figure S2.** Rootbox system, cultivation conditions, and imaging platforms used in this study.

**(A, B)** Custom-made rootbox equipped with oxygen-sensitive optode foil. The **(A)** front and **(B)** side views are shown.

**(C)** Rice plants cultivated in a bucket under controlled conditions in a growth chamber.

**(D–F)** Planar optode imaging setup in a dark room. **(D)** Overview of the imaging platform and **(E)** a close-up view are shown. The fluorescence emission from the optode foil induced by the blue LED excitation is shown in panel **(F)**. The position of the optode foil is indicated by dashed lines in panels **(E)** and **(F)**.

**(G, H)** X-ray CT scanning system of the rootboxes for non-destructive root imaging. The major components are as follows: (a) an X-ray source tube, (b) a turntable, and (c) a detector.

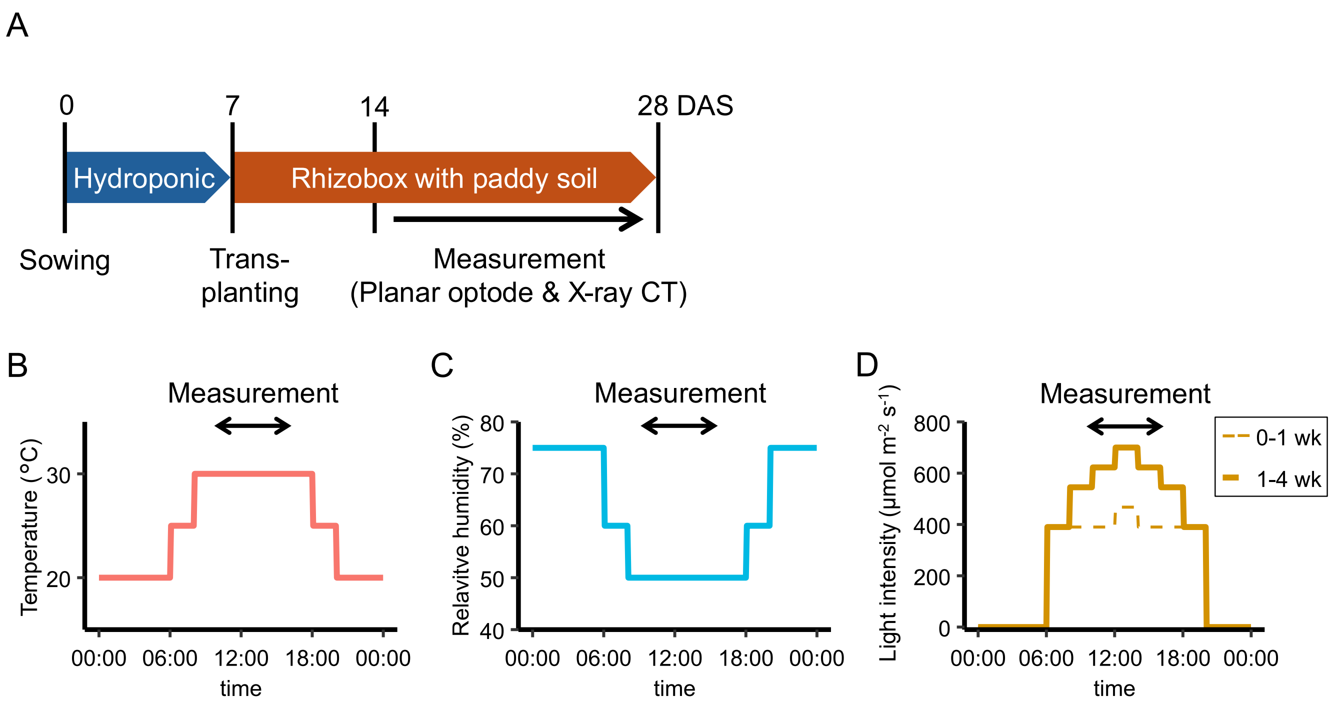


**Figure S3.** Cultivation conditions used in this study.

**(A)** Schematic illustration of cultivation schedule. DAS: days after sowing.

**(B–D)** Daily programmed changes in the **(B)** air temperature, **(C)** relative humidity, and **(D)** light intensity in the growth chamber.

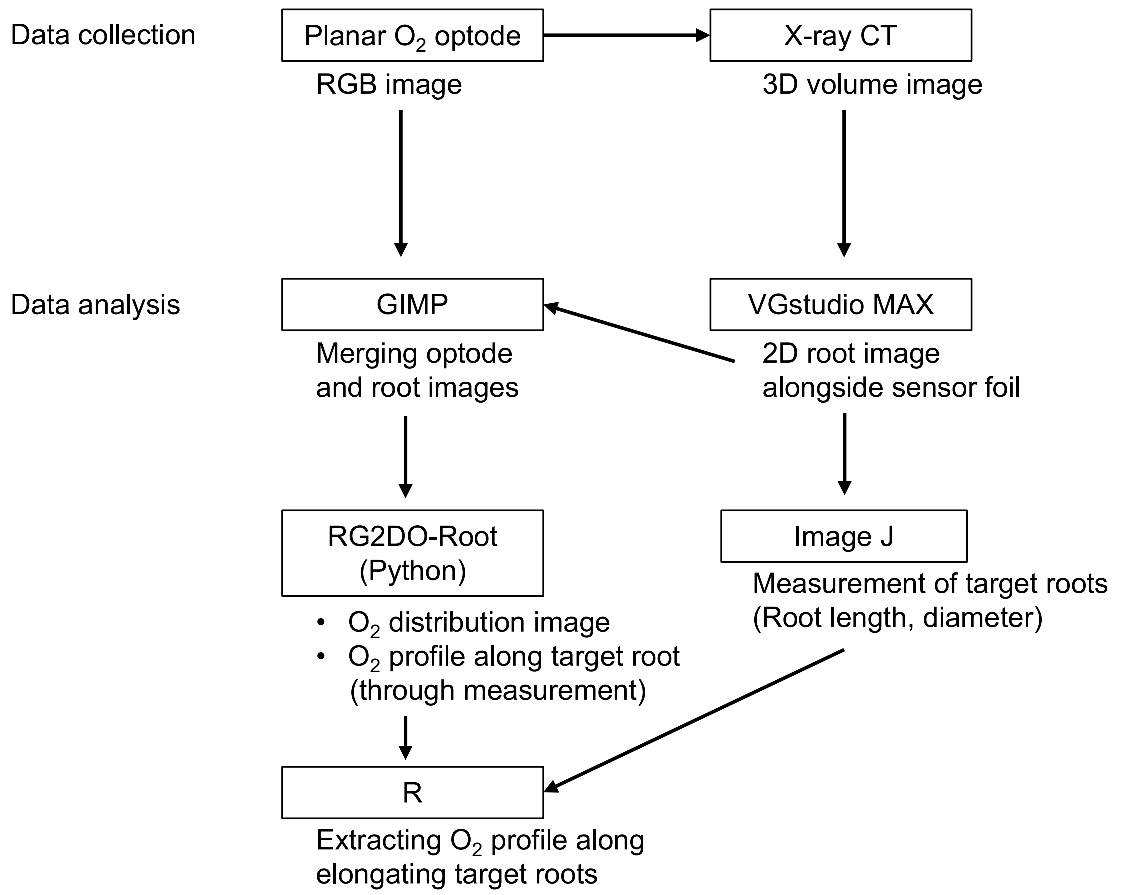


**Figure S4.** Workflow for multimodal data collection and analysis of rhizosphere dissolved oxygen (DO) distribution and root development.

Planar oxygen optode imaging was used to obtain RGB images of rhizosphere DO distributions, whereas X-ray CT was applied to acquire 3D root volume images. 2D root images adjacent to the optode sensor foil were extracted from CT data using VGStudio MAX. Optode and root images were merged using GIMP, and DO distribution images and longitudinal DO profiles along target roots were quantified using a custom Python-based tool (RG2DO-Root). Root traits, including length and diameter, were measured from CT images using ImageJ software. Finally, temporal changes in oxygen profiles along elongating target roots were analyzed using R.

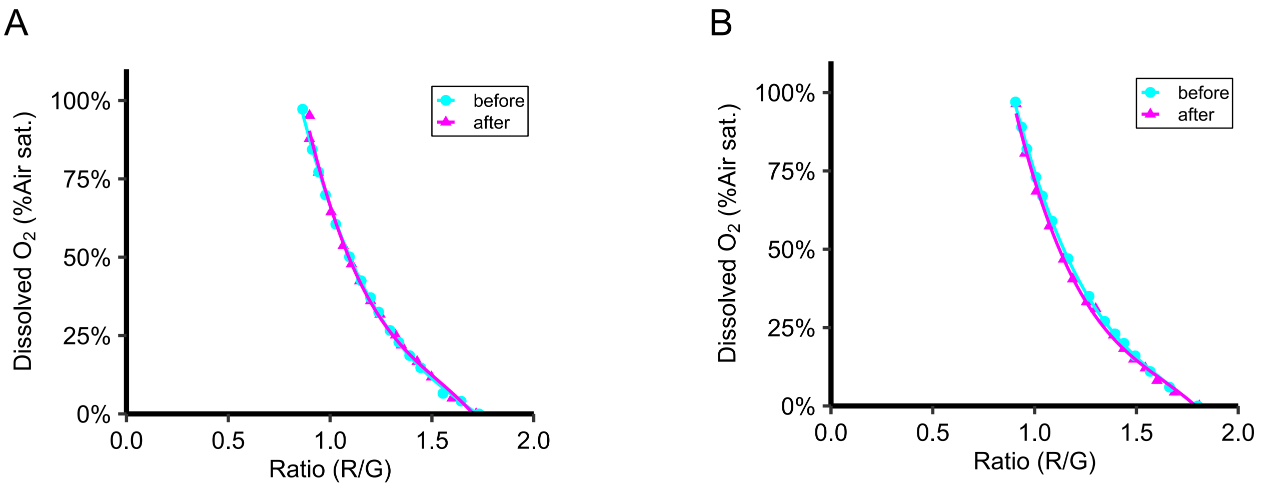


(before) $y = -1.04x^3 + 5.22x^2 - 9.19x + 5.68$
$R^2 = 0.999$, $P = 1.50e\text{-}19^{**}$

(after) $y = -1.76x^3 + 8.06x^2 - 12.88x + 7.25$
$R^2 = 0.996$, $P = 1.11e\text{-}13^{**}$

Model comparison: $P = 0.24$

(before) $y = -1.10x^3 + 5.54x^2 - 9.82x + 6.13$
$R^2 = 1.00$, $P = 7.36e\text{-}20^{**}$

(after) $y = -1.38x^3 + 6.74x^2 - 11.44x + 6.80$
$R^2 = 0.998$, $P = 1.27e\text{-}14^{**}$

Model comparison: $P = 0.16$

**Figure S5.** Calibration of oxygen-sensitive optode foils before and after measurements.
Representative calibration curves obtained from two optode foils are shown. In each panel, different colors indicate the timing of calibration measurements. For each time point, dissolved oxygen (DO) profiles were fitted using a third-order polynomial regression model (DO ~ ratio³). Solid lines represent fitted regression curves. Coefficients of determination ($R^2$) are shown in each panel. Statistical significance of the fitted polynomial regression models was evaluated using *F*-tests ($^{**}P < 0.01$). Differences in curve shapes among time points were assessed by comparing models with and without interaction terms between ratio and time.

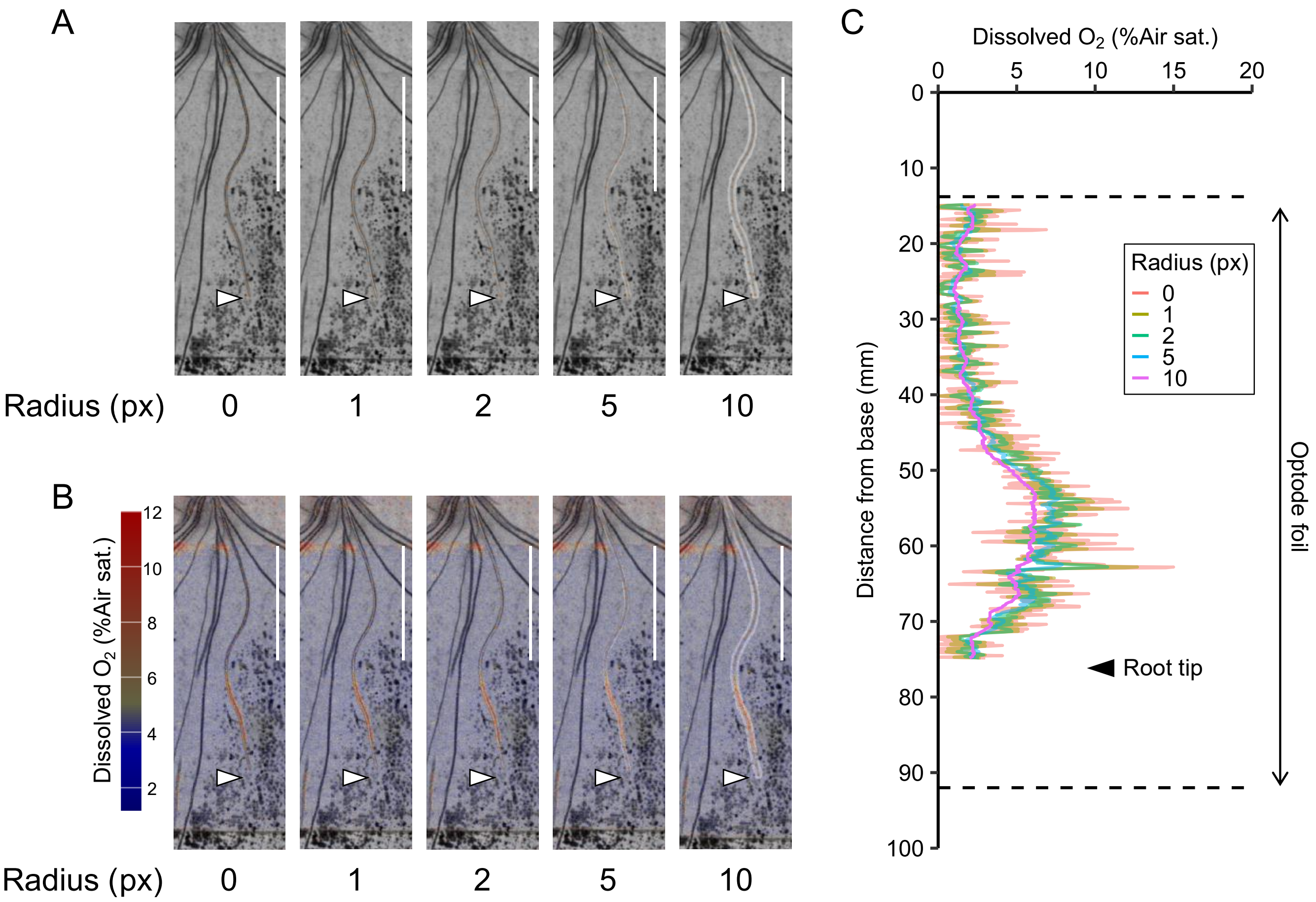


**Figure S6.** Comparison of dissolved oxygen (DO) profiles along a crown root obtained using different circular averaging radii in 'Koshihikari' (KSH) at 24 days after sowing.

**(A)** Traced crown root images using different circular averaging radii. Scale bars = 3 cm.

**(B)** Corresponding DO distribution images in panel **(A)**. Colors indicate DO concentrations.

**(C)** Longitudinal DO profiles along the crown root for each averaging radius. Dashed lines indicate optode foil upper and lower boundaries.

Arrowheads indicate root tip position.

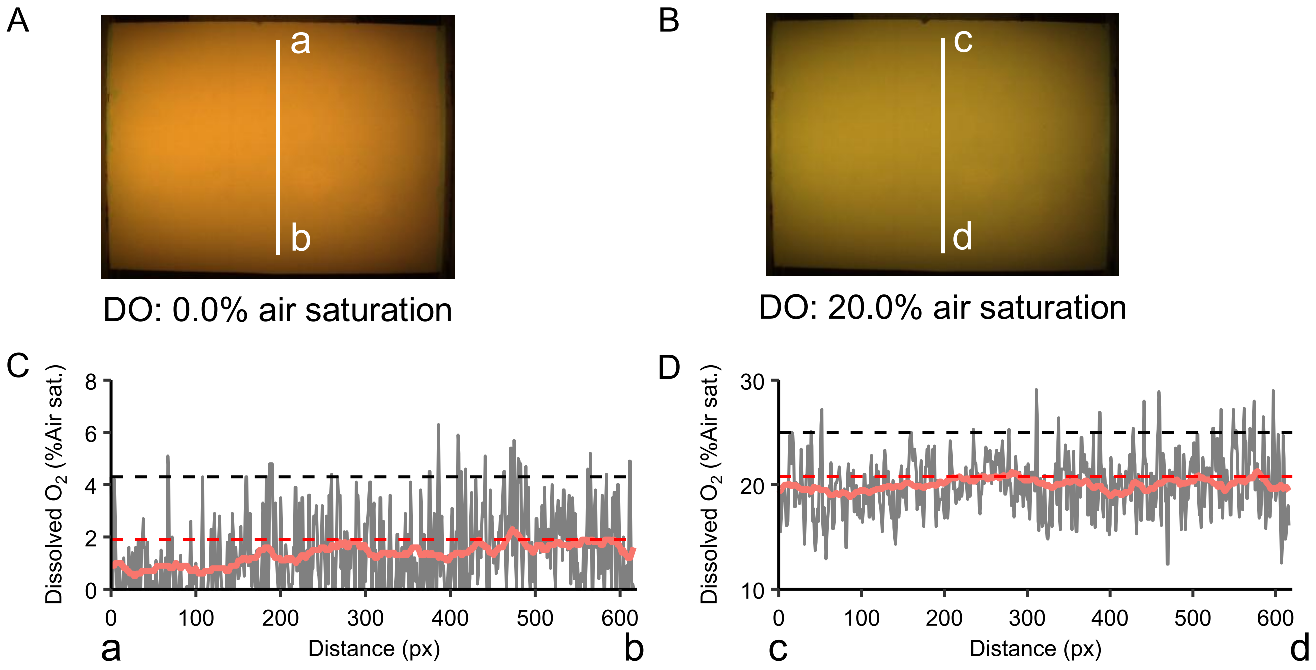


E

Integration radius = 0 px

| Calibration DO (%) | Measured DO by RG2DO-Root (%) | SD (%) | Upper noise boundary (95th percentile, %) |
|---|---|---|---|
| 0.0 | 1.4 | 1.5 | 4.3 |
| 3.8 | 3.8 | 1.8 | 6.9 |
| 11.3 | 12.0 | 2.3 | 15.6 |
| 14.2 | 14.6 | 2.4 | 18.4 |
| 16.5 | 16.7 | 2.6 | 21.1 |
| 20.0 | 20.0 | 2.9 | 25.0 |

F

Integration radius = 10 px

| Calibration DO (%) | Measured DO by RG2DO-Root (%) | SD (%) | Upper noise boundary (95th percentile, %) |
|---|---|---|---|
| 0.0 | 1.3 | 0.4 | 1.9 |
| 3.8 | 3.6 | 0.4 | 4.2 |
| 11.3 | 11.8 | 0.5 | 12.7 |
| 14.2 | 14.2 | 0.5 | 15.0 |
| 16.5 | 16.5 | 0.4 | 17.2 |
| 20.0 | 20.0 | 0.5 | 20.8 |

**Figure S7.** Sensor noise characteristics of planar optodes and noise reduction by spatial averaging.

**(A, B)** RGB images of oxygen-sensitive optode foils exposed to calibration solutions with different dissolved oxygen (DO) levels.

**(C, D)** DO profiles along the central vertical line in panels **(A)** and **(B)**. DO values obtained using integration radii of 0 px (gray line) and 10 px (red line) are presented for each DO level. Letters at the ends of the x-axis indicate the corresponding positions in panels **(A)** and **(B)**. Dashed lines indicate the upper noise boundary for each averaging radius.

**(E, F)** DO values measured using RG2DO-Root and corresponding upper noise boundaries across calibration DO levels for integration radii of **(E)** 0 px and **(F)** 10 px.

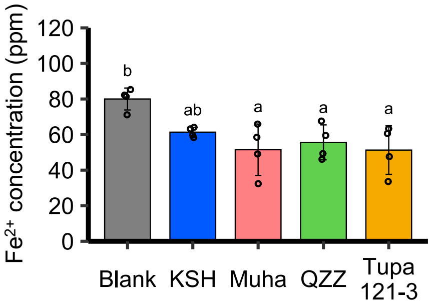


**Figure S8.** Ferrous iron concentrations in the soil pore water of a blank rootbox (without plants) and rootboxes planted with different rice cultivars at 20 days after sowing.

Bars represent mean ± SD (*n* = 4 samples). Different letters indicate significant differences among cultivars ($P$ < 0.05, Tukey's multiple comparison test).

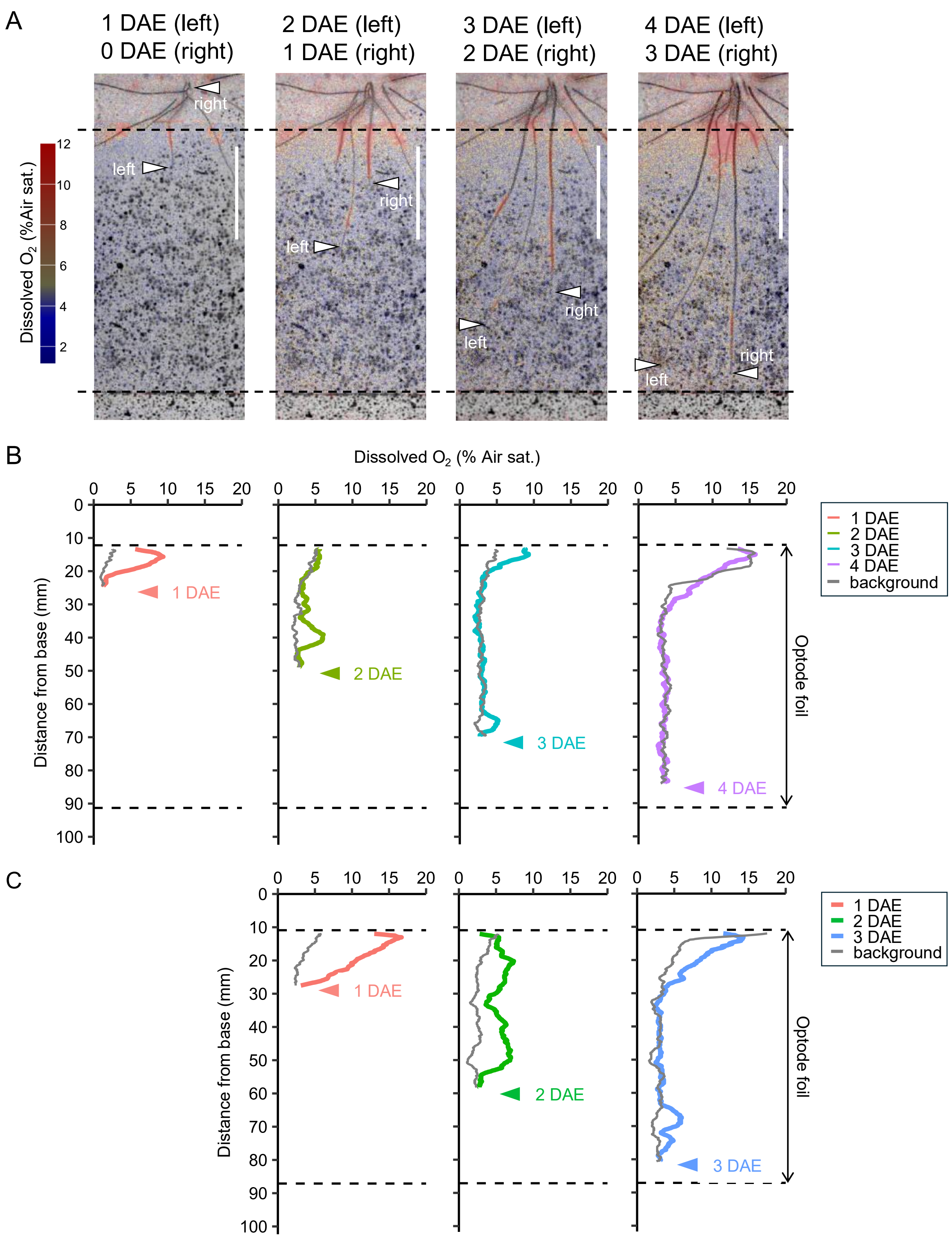


**Figure S9.** Daily changes in longitudinal dissolved oxygen (DO) profiles along two crown roots of KSH at 18–21 days after sowing.

**(A)** DO concentrations along the crown roots. Arrowheads indicate root tip positions. Colors represent DO concentrations. Scale bars = 3 cm.

**(B, C)** Longitudinal DO profiles for **(B)** left and **(C)** right roots in panel **(A)**. Colored lines indicate rhizosphere DO values, whereas gray lines represent background DO levels.

Dashed lines denote optode foil upper and lower boundaries. Arrowheads indicate root tip positions. DAE: days after emergence.

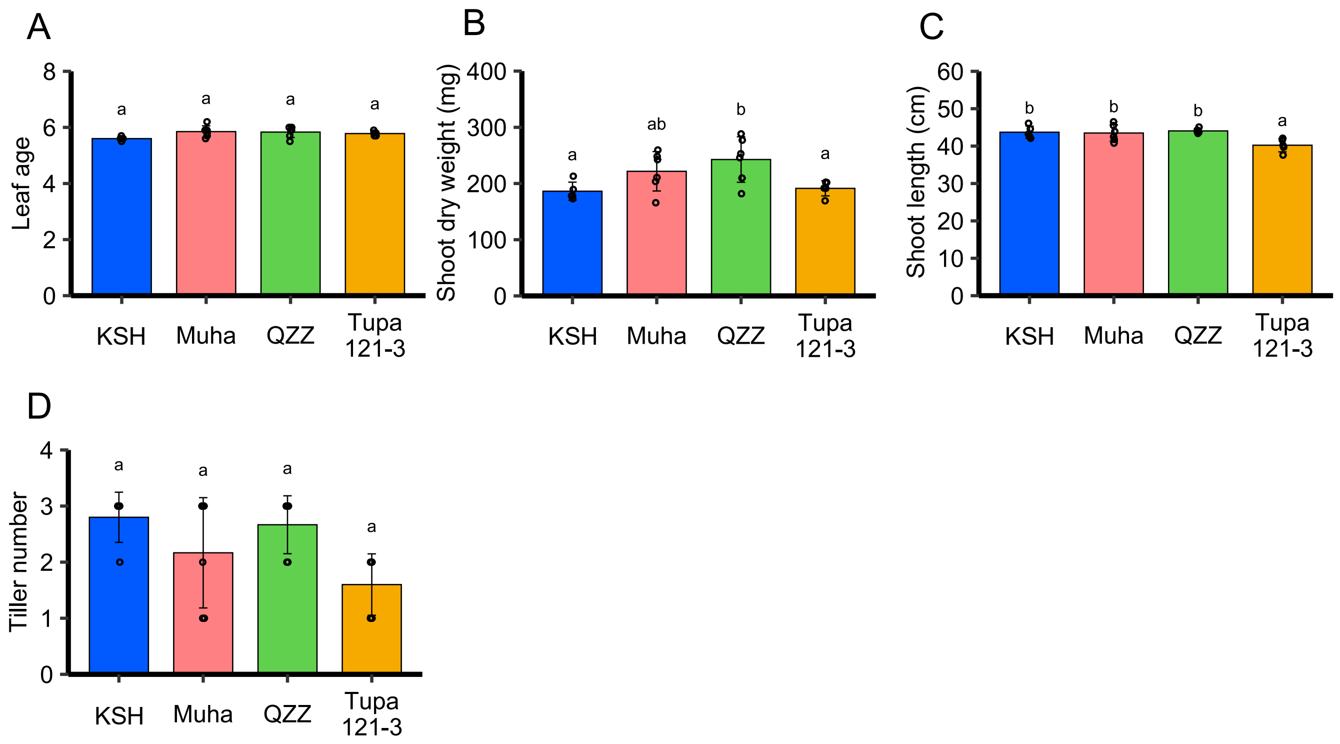


**Figure S10.** Shoot traits of four rice cultivars at 21 days after sowing.

**(A–D)** Quantification of shoot traits in each cultivar. Bars represent mean ± SD (*n* = 5 or 6 plants). Different letters indicate significant differences among cultivars ($P < 0.05$, Tukey's multiple comparison test).

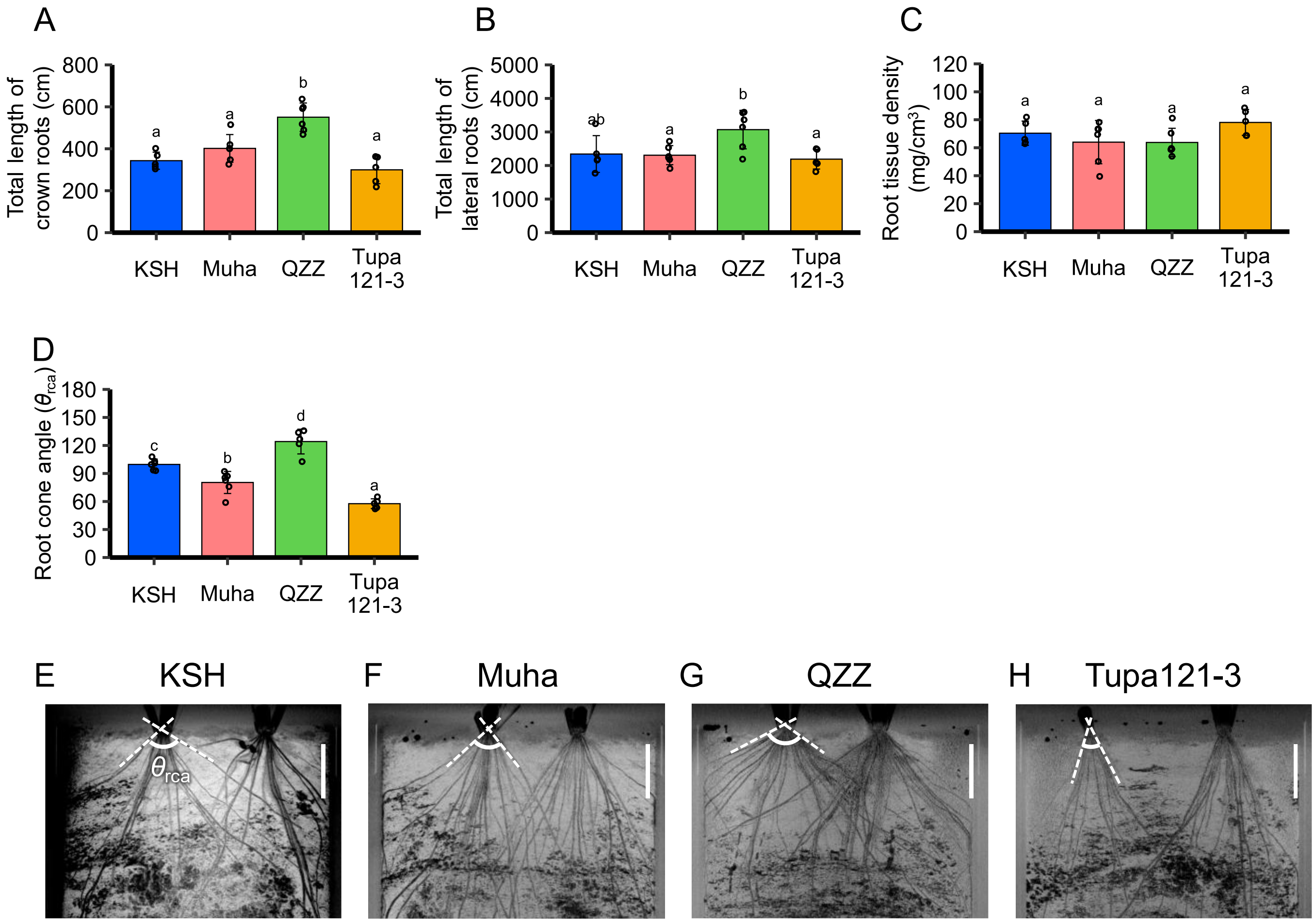


**Figure S11.** Root traits of four rice cultivars at 21 days after sowing.

**(A–D)** Quantification of root traits in each cultivar. Bars represent mean ± SD ($n$ = 5 or 6 plants). Different letters indicate significant differences among cultivars ($P$ < 0.05, Tukey's multiple comparison test).

**(E–H)** Root system architecture visualized via X-ray CT for each cultivar. $\theta_{rca}$ is the root cone angle. Scale bar = 2 cm.

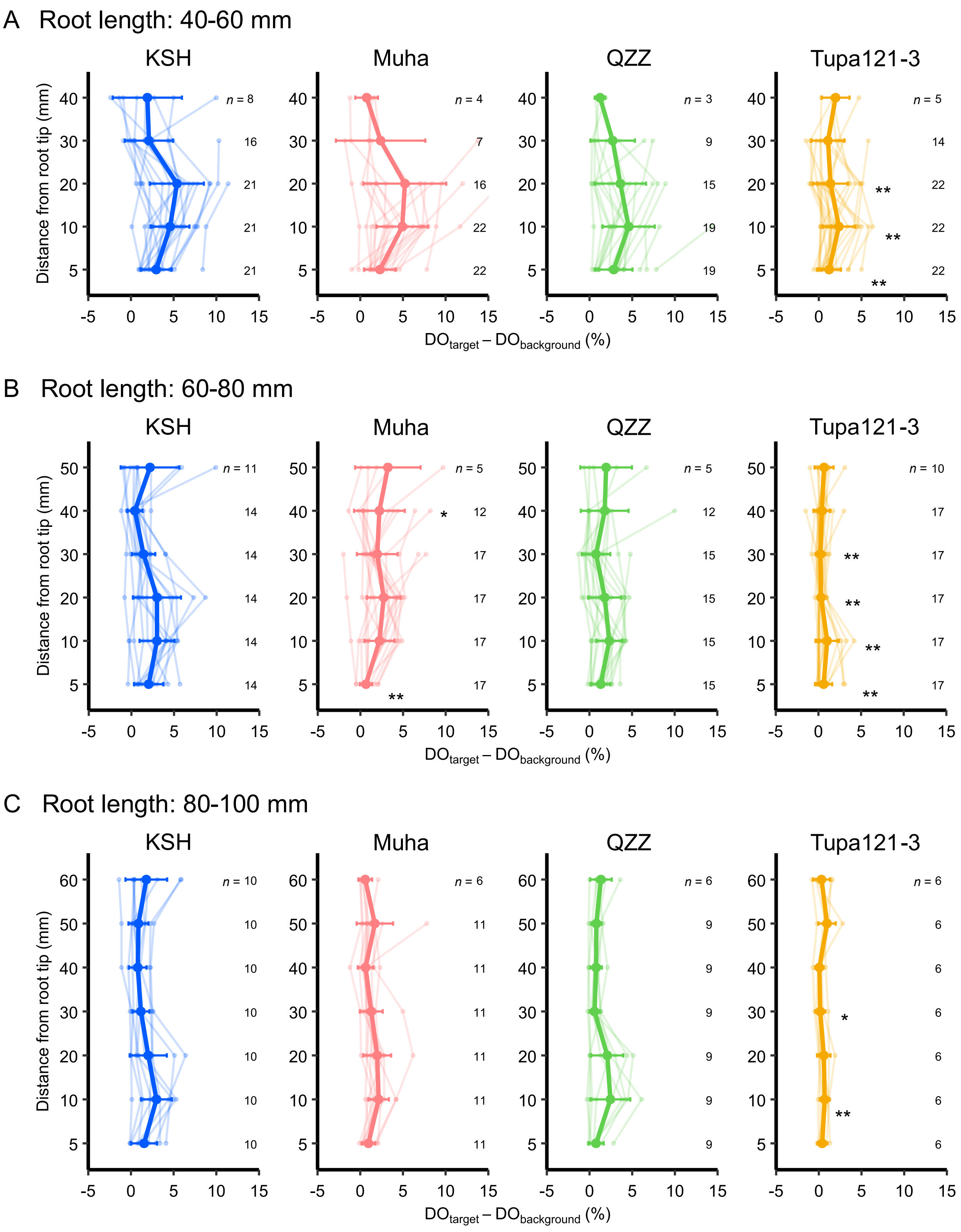


**Figure S12.** Longitudinal profiles of differences in dissolved oxygen (DO) levels between the rhizosphere and background along crown roots across different root-length classes for four rice cultivars.

Asterisks denote significant differences between each cultivar and KSH (two-tailed Student's *t* test; **P* < 0.05, ***P* < 0.01). The numbers adjacent to the data points indicate the number of replicates.

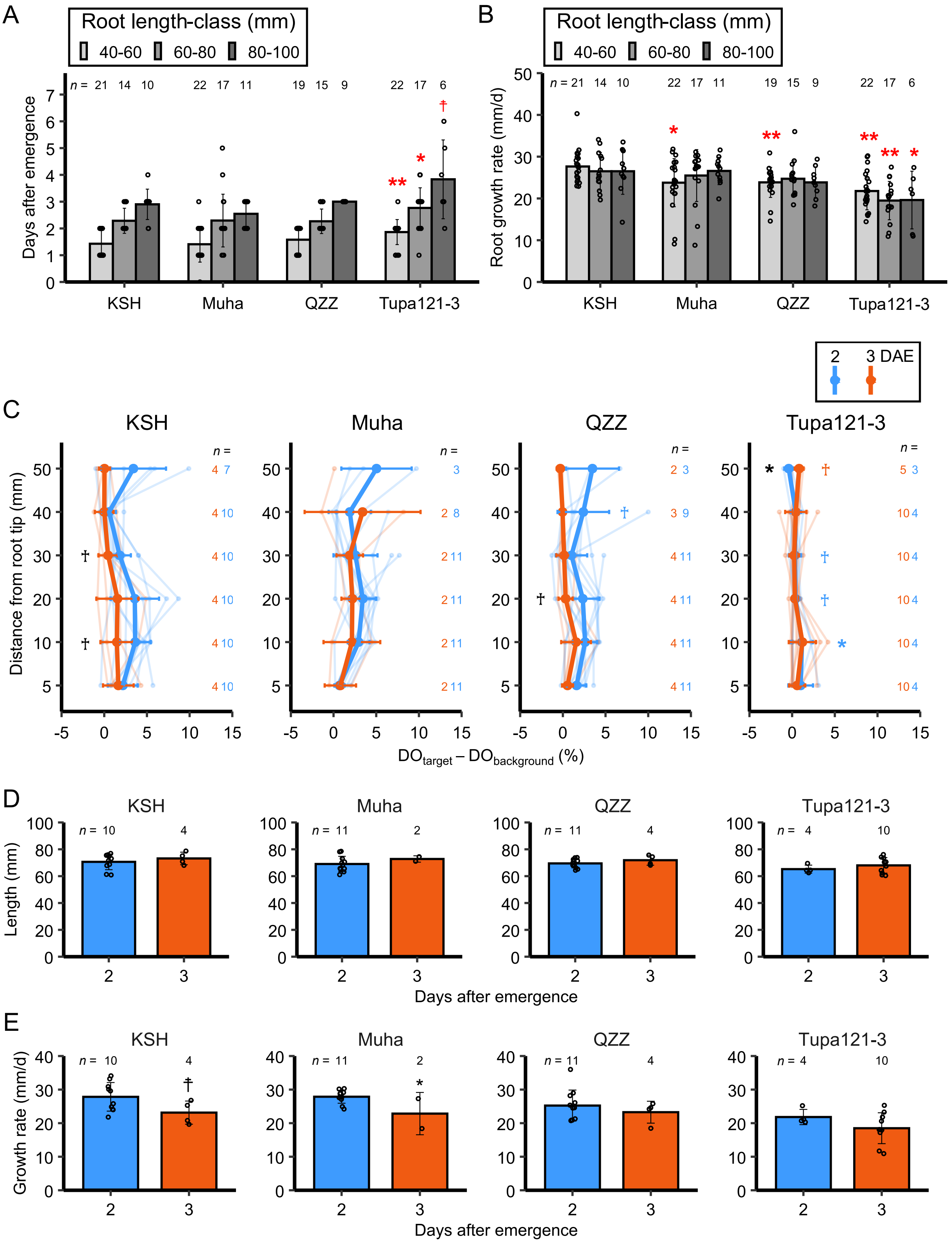


**Figure S13.** Effects of root age on dissolved oxygen (DO) profiles in four rice cultivars.

**(A)** Root age and **(B)** growth rate across different root-length classes for each cultivar. Bars represent mean ± SD. Asterisks denote significant differences between each cultivar and KSH for each root-length class (two-tailed Student's *t* test; †*P* < 0.1, **P* < 0.05, ***P* < 0.01). The numbers above the data points indicate the number of replicates.

**(C)** Longitudinal profiles of differences in DO levels along crown roots with lengths of 60–80 mm at different root ages for the four cultivars. Asterisks on the right denote significant differences between each cultivar and KSH for each root age (two-tailed Student's *t* test; †*P* < 0.1, **P* < 0.05, ***P* < 0.01). Asterisks on the left denote significant differences between rhizosphere and background values for each cultivar. Numbers adjacent to the data points indicate the number of replicates for each root age. Colors indicate the root age (DAE, days after emergence). **(D)** Root length and **(E)** growth rate at different root ages for crown roots with lengths of 60–80 mm for the four cultivars. Asterisks denote significant differences between 2 and 3 DAE (two-tailed Student's *t* test; †*P* < 0.1, **P* < 0.05, ***P* < 0.01).

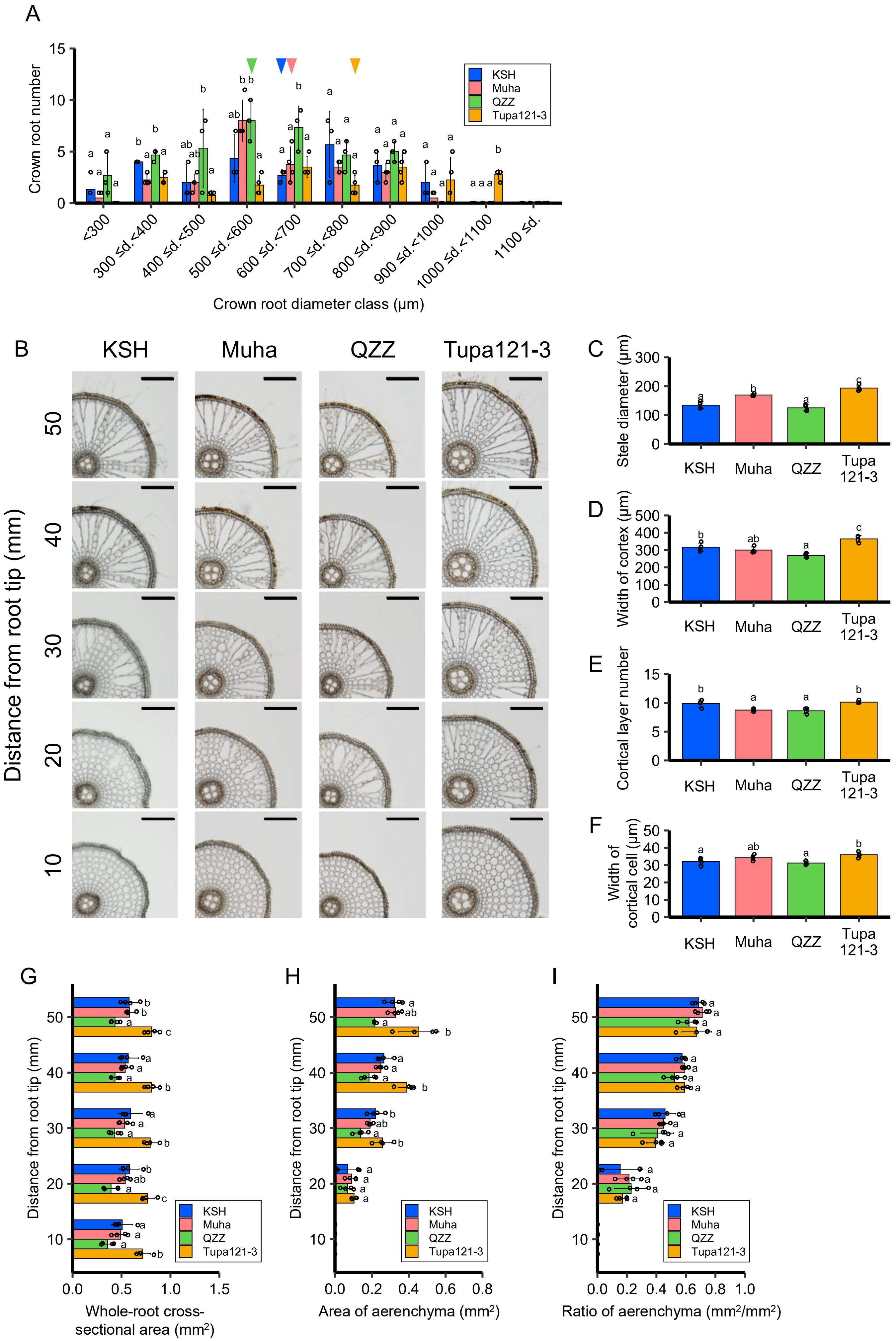

A
Crown root number
KSH
Muha
QZZ
Tupa121-3
<300
300 ≤d.<400
400 ≤d.<500
500 ≤d.<600
600 ≤d.<700
700 ≤d.<800
800 ≤d.<900
900 ≤d.<1000
1000 ≤d.<1100
1100 ≤d.
Crown root diameter class (μm)
B
KSH
Muha
QZZ
Tupa121-3
Distance from root tip (mm)
50
40
30
20
10
C
Stele diameter (μm)
D
Width of cortex (μm)
E
Cortical layer number
F
Width of cortical cell (μm)
KSH
Muha
QZZ
Tupa 121-3
G
Distance from root tip (mm)
Whole-root cross-sectional area (mm2)
H
Area of aerenchyma (mm2)
I
Ratio of aerenchyma (mm2/mm2)

**Figure S14.** Anatomical structure of crown roots in four rice cultivars at 21 days after sowing.

**(A)** Distribution of crown root diameters in each cultivar. Rectangles indicate mean diameters. Bars represent mean ± SD ($n$ = 3 or 4 plants). Different letters indicate significant differences among cultivars within each diameter class ($P < 0.05$, Tukey's multiple comparison test). d.: diameter.

**(B)** Representative cross-sectional images of crown roots at different distances from the root tip for each cultivar. Scale bars = 200 μm.

**(C–F)** Quantification of root anatomical traits at 50 mm from the root tip for each cultivar. Bar plots represent mean ± SD ($n$ = 4 roots). Different letters indicate significant differences among cultivars ($P < 0.05$, Tukey's multiple comparison test).

**(G–I)** Quantification of total root cross-sectional area and aerenchyma formation in panel **(B)**.

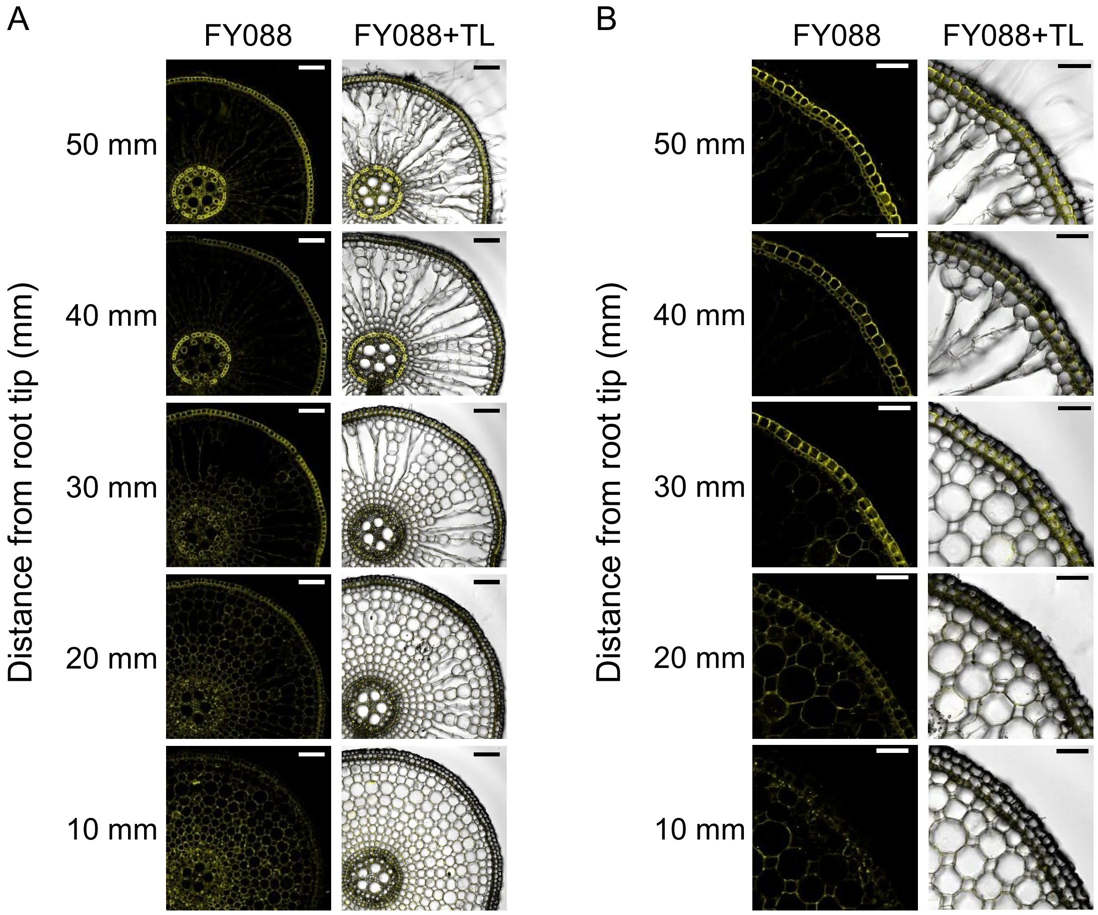


**Figure S15.** Suberin lamella formation at different distances from the root tip in a crown root of 'Tupa121-3'. Fluorol Yellow 088 (FY088)-stained root sections and corresponding merged transmitted light (TL) images. **(A)** Low-magnification overview images and **(B)** enlarged images of the outer cell layers. Scale bars = **(A)** 100 µm and **(B)** 50 µm.
