## Supplementary material for "Characterization of Rhizosphere Oxidation Associated with Root Development in Rice Using Planar Oxygen Optodes": RG2DO-Root: __main__.py

```
from PySide6.QtWidgets import QApplication

from modules.gui import QtMainWindow


def main():
    app = QApplication()
    main_wind = QtMainWindow()
    main_wind.show()

    app.exec()


if __name__ == "__main__":
    main()
```
