## Supplementary material for "Characterization of Rhizosphere Oxidation Associated with Root Development in Rice Using Planar Oxygen Optodes": RG2DO-Root: coordinates.py

```
from __future__ import annotations

from dataclasses import dataclass
from functools import cache

import numpy as np
import polars as pl
from skimage.morphology import disk


@dataclass
class Df_for_Drawing:
    y_ary: np.ndarray
    x_ary: np.ndarray
    color: tuple[int]

    def __post_init__(self):
        self.y_ary = np.asarray(self.y_ary)
        self.x_ary = np.asarray(self.x_ary)

        assert self.y_ary.shape == self.x_ary.shape

    def __getitem__(self, item):
        if isinstance(item, np.ndarray):
            return Df_for_Drawing(
                self.y_ary[item],
                self.x_ary[item],
                self.color,
            )

        assert NotImplementedError()


@cache
def get_ofset_list(radius: int):
    b = disk(radius=radius)
    return np.array(np.where(b == 1)).transpose() - radius


def remove_duplicates(np_ary: np.ndarray, img_shape: tuple[int, int]):
    df = pl.DataFrame(
        {
            "y": np_ary[0],
            "x": np_ary[1],
        }
    )
    df = (
        df.unique(subset=["y", "x"])
        .filter(pl.col("y") >= 0)
        .filter(pl.col("x") >= 0)
        .filter(pl.col("y") < img_shape[0])
        .filter(pl.col("x") < img_shape[1])
    )
    return df.to_numpy().transpose()


def get_df_for_drawing(
    y_ary: np.ndarray,
    x_ary: np.ndarray,
    radius: int,
    img_shape: tuple[int, int],
    color: tuple[int] = (0, 0, 0, 0),
):
    yx_ary = np.array([y_ary, x_ary], dtype=np.int16)

    res_yx_ary_list = []
    for size in [radius]:
        sub_yx_ary = yx_ary
        sub_yx_ary = np.expand_dims(sub_yx_ary, 0)
        offset_list = get_ofset_list(size)
        offset_list = np.expand_dims(offset_list, 2)
        res_yx_ary = sub_yx_ary + offset_list
        res_yx_ary = np.concatenate(res_yx_ary, axis=1)
        res_yx_ary = remove_duplicates(res_yx_ary, img_shape)
        res_yx_ary_list.append(res_yx_ary)

    res_yx_ary_ary = np.concatenate(res_yx_ary_list, axis=1)
    res_yx_ary_ary = remove_duplicates(res_yx_ary_ary, img_shape)

    return Df_for_Drawing(*res_yx_ary_ary, color=color)
```
