## Supplementary material for "Characterization of Rhizosphere Oxidation Associated with Root Development in Rice Using Planar Oxygen Optodes": RG2DO-Root: gui.py

```
from __future__ import annotations

import os
import pickle
import re
from dataclasses import dataclass, field
from pathlib import Path

import numpy as np
import polars as pl
import PySide6  # NOQA
from PySide6.QtCore import Qt
from PySide6.QtGui import QAction, QDragEnterEvent, QDropEvent
from PySide6.QtWidgets import QMainWindow, QMenuBar
from scipy import interpolate
from skimage import io

from .coordinates import get_df_for_drawing
from .toolbar import QtToolBar

if True:
    import pyqtgraph as pg  # NOQA
    from pyqtgraph import CONFIG_OPTIONS, mkColor

    CONFIG_OPTIONS["imageAxisOrder"] = "row-major"


def rgb2o(np_ary: np.ndarray):
    np_ary = np_ary.astype(np.float32)
    r = np_ary[..., 0]  # noqa
    g = np_ary[..., 1]  # noqa
    b = np_ary[..., 2]  # noqa
    np_ary = -132.3 * (r / (g + 0.0001)) + 201.0
    np_ary = np_ary.clip(0.0, 100.0)
    return np_ary


@dataclass
class Coordinate:
    x: int
    y: int

    def get_distance_to(self, target: Coordinate):
        return ((self.x - target.x) ** 2 + (self.y - target.y) ** 2) ** 0.5


@dataclass
class PathPaser:
    src_img_path: Path

    @property
    def _re(self):
        return re.findall("(.*)_(.*)_(.*).png", self.src_img_path.name)

    @property
    def name(self):
        return self._re[0][0]

    @property
    def type(self):
        return self._re[0][1].lower()

    @property
    def suffix(self):
        return self._re[0][2]


@dataclass
class ListForDf:
    root_id: list[int] = field(default_factory=list)
    distance: list[float] = field(default_factory=list)
    y: list[float] = field(default_factory=list)
    x: list[float] = field(default_factory=list)
    o2: list[str] = field(default_factory=list)
    count_all: list[int] = field(default_factory=list)
    count_gt1: list[int] = field(default_factory=list)
    suffix: list[str] = field(default_factory=list)

    def append(self, root_id, distance, y, x, o2, count_all, count_gt1, suffix):
        self.root_id.append(root_id)
        self.distance.append(distance)
        self.y.append(y)
        self.x.append(x)
        self.count_all.append(count_all)
        self.count_gt1.append(count_gt1)
        self.o2.append(f"{o2:0.1f}")
        self.suffix.append(suffix)


class Polyline:
    def __init__(self) -> None:
        self.__pos_list: list[Coordinate] = []
        self.__current = 0

    def append(self, pos: Coordinate):
        self.__pos_list.append(pos)

    def delete(self, i: int):
        del self.__pos_list[i]

    @property
    def x_list(self):
        return [c.x for c in self]

    @property
    def y_list(self):
        return [c.y for c in self]

    @property
    def distance_list(self):
        distance_list = [0.0]
        for pos_i, pos in enumerate(self):
            if pos_i:
                distance_list.append(pos.get_distance_to(self[pos_i - 1]))

        return distance_list

    def __getitem__(self, key):
        return self.__pos_list[key]

    def __len__(self):
        return len(self.__pos_list)

    def __iter__(self):
        return self

    def __next__(self):
        if self.__current == len(self.__pos_list):
            self.__current = 0
            raise StopIteration()

        self.__current += 1
        return self.__pos_list[self.__current - 1]


def get_df(img_path: Path, polyline_list: list[Polyline], size: int):
    img = io.imread(img_path)
    pp = PathPaser(img_path)

    list_df = ListForDf()
    for pl_i, polyline in enumerate(polyline_list):
        distance_list = polyline.distance_list
        step_list = np.arange(0, np.sum(distance_list), 1)

        y_interpolate = interpolate.interp1d(np.cumsum(distance_list), polyline.y_list)(
            step_list
        )

        x_interpolate = interpolate.interp1d(np.cumsum(distance_list), polyline.x_list)(
            step_list
        )

        for step, y, x in zip(step_list, y_interpolate, x_interpolate, strict=True):
            pos = get_df_for_drawing(
                np.asarray([np.ceil(y).astype(np.uint16)]),
                np.asarray([np.ceil(x).astype(np.uint16)]),
                size,
                img.shape[0:2],
            )
            o2 = rgb2o(img[pos.y_ary, pos.x_ary, :])
            count_all = len(o2)
            count_gt1 = np.sum(o2 > 1.0)
            o2 = o2.mean()
            list_df.append(pl_i, step, y, x, o2, count_all, count_gt1, pp.suffix)

    return pl.DataFrame(list_df.__dict__)


class QtMenubar(QMenuBar):
    def __init__(self, parent: QtMainWindow, *args, **kwargs):
        super().__init__(parent, *args, **kwargs)
        self.__parent = parent
        self.build()

    def build(self):
        self.act_save = QAction(
            text="save",
            parent=self.parent(),
            shortcut="Ctrl+S",
            triggered=self.on_act_save,
        )
        self.addAction(self.act_save)

        self.act_load = QAction(
            text="load",
            parent=self.parent(),
            shortcut="Ctrl+L",
            triggered=self.on_act_load,
        )
        self.addAction(self.act_load)

        self.act_export = QAction(
            text="export",
            parent=self.parent(),
            shortcut="Ctrl+R",
            triggered=self.on_act_export,
        )
        self.addAction(self.act_export)

    def on_act_save(self):
        iv = self.__parent.image_view
        save_dict = {"polyline list": iv.polyline_list}

        with open(self.__parent.pkl_path, "wb") as f:
            pickle.dump(save_dict, f)

    def on_act_load(self):
        if not self.__parent.pkl_path.is_file():
            return

        iv = self.__parent.image_view
        with open(self.__parent.pkl_path, "rb") as f:
            load_dict = pickle.load(f)

        iv.polyline_list = load_dict["polyline list"]
        if len(iv.polyline_list):
            iv.current_line = 0
        else:
            iv.current_line = -1

        iv.update_marks()

    def on_act_export(self):
        main = self.__parent
        iv = main.image_view
        polyline_list = iv.polyline_list

        df_list = []
        for img_path in main.po2o_image_paths:
            df_list.append(
                get_df(
                    img_path,
                    polyline_list,
                    self.__parent.toolbar.spinbox_radius.value(),
                )
            )

        df = pl.concat(df_list)
        df.write_csv(main.csv_path)


class QtMainWindow(QMainWindow):
    def __init__(self, **kwargs) -> None:
        super().__init__(**kwargs)

        self.init_interface()
        self.load_test_images()

        self.setAcceptDrops(True)

    def init_interface(self):
        self.resize(800, 800)
        self.setWindowTitle("Planar O2 Optode")

        self.toolbar = QtToolBar(self)
        self.addToolBar(self.toolbar)

        self.image_view = QtImageView(parent=self)
        self.setCentralWidget(self.image_view)

        self.menubar = QtMenubar(self)
        self.setMenuBar(self.menubar)

        self.image_view.update_img_item_o2()

    def dragEnterEvent(self, ev: QDragEnterEvent):
        ev.accept()

    def dropEvent(self, ev: QDropEvent):
        ev.accept()

        flist = [u.toLocalFile() for u in ev.mimeData().urls()]
        if len(flist) != 1:
            return

        self.src_dir = Path(flist[0]).resolve()
        self.load_images(self.src_dir)
        self.image_view.reset_polyline()

    def load_images(self, src_dir: Path):
        img_file_paths = [
            Path(src_dir, f)
            for f in sorted(os.listdir(src_dir))
            if f.lower().endswith(".png")
        ]
        self.ct_image_path = Path("")
        self.po2o_image_paths: list[Path] = []
        for f in img_file_paths:
            pp = PathPaser(f)
            if pp.type == "ct":
                self.ct_image_path = f
            elif pp.type == "rgb":
                self.po2o_image_paths.append(f)

        self.ct_image = io.imread(self.ct_image_path)

        self.image_view.img_item_ct.setImage(self.ct_image)

        self.o2_images = []
        for p in self.po2o_image_paths:
            img = io.imread(p)
            img = rgb2o(img)
            self.o2_images.append(img)

        self.image_view.img_item_o2.setImage(self.o2_images[0])

    def load_test_images(self):
        self.src_dir = Path("target_data").resolve()
        self.load_images(self.src_dir)

    @property
    def pkl_path(self):
        return Path(str(self.src_dir) + ".pkl")

    @property
    def csv_path(self):
        return Path(str(self.src_dir) + ".csv")


class QtImageView(pg.GraphicsLayoutWidget):
    def __init__(self, parent: QtMainWindow):
        super().__init__(parent=parent)
        self.__main_window = parent
        self.__toolbar = self.__main_window.toolbar

        self.view_box = pg.ViewBox(lockAspect=True, invertY=True)
        self.img_item_ct = pg.ImageItem()
        self.view_box.addItem(self.img_item_ct)
        self.img_item_o2 = pg.ImageItem()
        self.img_item_range = pg.ImageItem()

        cm = pg.ColorMap(
            pos=(0.0, 10.0 / 100.0, 20.0 / 100.0, 30.0 / 100.0, 1.0),
            color=(
                mkColor("#0000FF00"),
                mkColor("#0000FF40"),
                mkColor("#FFFF0040"),
                mkColor("#FF000040"),
                mkColor("#FF000040"),
            ),
        )
        self.img_item_o2.setColorMap(cm)
        bar = pg.ColorBarItem(values=(0, 30), colorMap=cm)
        bar.setImageItem(self.img_item_o2)
        self.view_box.addItem(self.img_item_range)
        self.view_box.addItem(self.img_item_o2)
        self.addItem(bar)
        self.addItem(self.view_box)

        self.marks = Marks(self)
        self.marks.show()
        self.view_box.addItem(self.marks)

        self.reset_polyline()

        self.img_item_ct.mouseClickEvent = self.on_mouse_clicked

    def update_img_item_o2(self):
        self.__toolbar.on_chkbox_oxygen_changed(
            self.__toolbar.chkbox_oxygen.checkState == Qt.CheckState.Checked
        )

    def reset_polyline(self):
        self.polyline_list: list[Polyline] = []
        self.current_line = -1
        self.update_marks()

    def on_mouse_clicked(self, ev):
        ev.accept()

        pos = Coordinate(int(ev.pos().x()), int(ev.pos().y()))
        if ev.button() == Qt.LeftButton:
            if ev.modifiers() & Qt.ShiftModifier:
                return
            if ev.modifiers() & Qt.ControlModifier:
                if self.current_line != -1:
                    pl = self.polyline_list[self.current_line]
                    pl.append(pos)
                    self.update_marks()
                return
            polyline = Polyline()
            polyline.append(pos)
            self.polyline_list.append(polyline)
            self.current_line = len(self.polyline_list) - 1
            self.update_marks()

    def update_marks(self):
        self.marks.draw(self.polyline_list, self.current_line)

        if self.img_item_ct.image is None:
            return

        radius = self.__main_window.toolbar.spinbox_radius.value()

        range_img = np.zeros(self.img_item_ct.image.shape, dtype=np.uint8)
        for _, polyline in enumerate(self.polyline_list):
            distance_list = polyline.distance_list
            step_list = np.arange(0, np.sum(distance_list), 1)

            y_interpolate = interpolate.interp1d(
                np.cumsum(distance_list), polyline.y_list
            )(step_list)

            x_interpolate = interpolate.interp1d(
                np.cumsum(distance_list), polyline.x_list
            )(step_list)

            y_interpolate = np.ceil(y_interpolate).astype(np.uint16)
            x_interpolate = np.ceil(x_interpolate).astype(np.uint16)

            if len(y_interpolate) != 0:
                if (
                    polyline.y_list[-1] != y_interpolate[-1]
                    or polyline.x_list[-1] != x_interpolate[-1]
                ):
                    y_interpolate = np.asarray(
                        y_interpolate.tolist() + [polyline.y_list[-1]]
                    ).astype(np.uint16)
                    x_interpolate = np.asarray(
                        x_interpolate.tolist() + [polyline.x_list[-1]]
                    ).astype(np.uint16)

            df = get_df_for_drawing(
                y_interpolate,
                x_interpolate,
                radius,
                range_img.shape[0:2],
                (255, 255, 255, 255),
            )
            range_img[df.y_ary, df.x_ary, :] = 255
            range_img[df.y_ary, df.x_ary, 3] = 64

        self.img_item_range.setImage(range_img)


class Marks(pg.GraphItem):
    def __init__(self, parent: QtImageView):
        self.__parent = parent
        self.dragPoint = None
        self.dragOffset = None
        self.textItems = []
        pg.GraphItem.__init__(self)
        self.scatter.sigClicked.connect(self.clicked)

    def draw(self, polyline_list: list[Polyline], current: int):
        if len(polyline_list) == 0:
            self.hide()
            return

        self.show()

        labels = []
        pos_list = []
        adj_list = []
        symbolPen_list = []
        symbolBrush_list = []
        lines_list = []
        self.pos_list_buf = []
        cnt = 0
        for ind, polyline in enumerate(polyline_list):
            if ind == current:
                symbol_pen = pg.mkPen((128, 64, 0))
                symbol_brush = pg.mkBrush((255, 64, 0, 64))
            else:
                symbol_pen = pg.mkPen((0, 128, 0))
                symbol_brush = pg.mkBrush((0, 255, 0, 64))
            for i, co in enumerate(polyline):
                if i == 0:
                    labels.append(str(ind))
                else:
                    labels.append("")
                pos_list.append([co.x, co.y])
                self.pos_list_buf.append([polyline, co, ind, i])
                symbolPen_list.append(symbol_pen)
                symbolBrush_list.append(symbol_brush)

                cnt += 1
                if i:
                    adj_list.append([cnt - 2, cnt - 1])
                    lines_list.append((255, 255, 255, 64, 3))

        pos = np.array(pos_list, dtype=float) + 0.5
        # adj = np.array(adj_list)
        self.setData(
            pos=pos,
            # adj=adj,
            symbolPen=symbolPen_list,
            symbolBrush=symbolBrush_list,
            size=5,
            # symbol="o",
            pxMode=False,
            text=labels,
            pen=np.array(
                lines_list,
                dtype=[
                    ("red", np.ubyte),
                    ("green", np.ubyte),
                    ("blue", np.ubyte),
                    ("alpha", np.ubyte),
                    ("width", float),
                ],
            ),
        )
        return

    def setData(self, **kwds):
        self.text = kwds.pop("text", [])
        self.data = kwds
        if "pos" in self.data:
            npts = self.data["pos"].shape[0]
            self.data["data"] = np.empty(npts, dtype=[("index", int)])
            self.data["data"]["index"] = np.arange(npts)
        self.setTexts(self.text)
        self.updateGraph()

    def setTexts(self, text):
        for i in self.textItems:
            i.scene().removeItem(i)
        self.textItems = []
        for t in text:
            item = pg.TextItem(t)
            self.textItems.append(item)
            item.setParentItem(self)

    def updateGraph(self):
        pg.GraphItem.setData(self, **self.data)
        for i, item in enumerate(self.textItems):
            item.setPos(*self.data["pos"][i])

    def mouseDragEvent(self, ev):
        if ev.button() != Qt.MouseButton.LeftButton:
            ev.ignore()
            return

        if ev.isStart():
            pos = ev.buttonDownPos()
            pts = self.scatter.pointsAt(pos)
            if len(pts) == 0:
                ev.ignore()
                return
            self.dragPoint = pts[0]
            ind = pts[0].data()[0]
            self.dragOffset = self.data["pos"][ind] - pos
        elif ev.isFinish():
            self.dragPoint = None
            return
        else:
            if self.dragPoint is None:
                ev.ignore()
                return

        ind = self.dragPoint.data()[0]
        pos = tuple(
            [int(i + j) for i, j in zip(ev.pos(), self.dragOffset, strict=True)]
        )
        self.pos_list_buf[ind][1].y = pos[1]
        self.pos_list_buf[ind][1].x = pos[0]
        self.data["pos"][ind] = pos
        self.updateGraph()
        ev.accept()

    def clicked(self, pts: pg.ScatterPlotItem, spot_items, ev):
        ind = spot_items[0].data()[0]
        if ev.modifiers() & Qt.ShiftModifier:
            polyline_list = self.__parent.polyline_list
            polyline = polyline_list[self.pos_list_buf[ind][2]]
            polyline.delete(self.pos_list_buf[ind][3])
            if len(polyline) == 0:
                del polyline_list[self.pos_list_buf[ind][2]]

                if len(polyline_list) == 0:
                    self.__parent.current_line = -1
                else:
                    self.__parent.current_line = 0

        else:
            self.__parent.current_line = self.pos_list_buf[ind][2]

        self.__parent.update_marks()
```
