## Supplementary material for "Characterization of Rhizosphere Oxidation Associated with Root Development in Rice Using Planar Oxygen Optodes": RG2DO-Root: toolbar.py

```
from __future__ import annotations

from typing import TYPE_CHECKING

from PySide6.QtCore import Qt
from PySide6.QtWidgets import QCheckBox, QLabel, QSpinBox, QToolBar

if TYPE_CHECKING:
    from .gui import QtMainWindow


class QtToolBar(QToolBar):
    def __init__(self, parent: QtMainWindow):
        super().__init__(*("Main",))
        self.main_window = parent
        self.setMovable(False)
        self.init_interface()

    def init_interface(self):
        self.chkbox_oxygen = QCheckBox(parent=self, text="Oxygen")
        self.chkbox_oxygen.stateChanged.connect(
            lambda: self.on_chkbox_oxygen_changed(
                self.chkbox_oxygen.checkState() == Qt.CheckState.Checked
            )
        )
        self.addWidget(self.chkbox_oxygen)

        self.addSeparator()
        self.addWidget(QLabel("Radius"))
        self.spinbox_radius = QtSpinBoxRadius(parent=self)
        self.spinbox_radius.valueChanged.connect(self.on_spinbox_changed)
        self.addWidget(self.spinbox_radius)

    def keyPressEvent(self, ev):
        ev.ignore()

    def keyReleaseEvent(self, ev):
        ev.ignore()

    def on_chkbox_oxygen_changed(self, state: bool):
        if state:
            self.main_window.image_view.img_item_o2.show()
        else:
            self.main_window.image_view.img_item_o2.hide()

    def on_spinbox_changed(self, val: int):
        self.main_window.image_view.update_marks()


class QtSpinBoxRadius(QSpinBox):
    def __init__(self, parent: QtToolBar, *args, **kwargs):
        super().__init__(parent, *args, **kwargs)
        self.toolbar = parent
        self.main_window = self.toolbar.main_window
        self.setToolTip("Radius [pixel]")
        self.setMaximum(10)
        self.setMinimum(0)
        self.setValue(1)
```
