## Supplementary material for "Characterization of Rhizosphere Oxidation Associated with Root Development in Rice Using Planar Oxygen Optodes": RG2DO-Root: pyproject.toml

```
[project]
name = "rg2do-root"
version = "0.1.0"
description = "Add your description here"
readme = "README.md"
requires-python = ">=3.14"
dependencies = [
    "numpy>=2.4.4",
    "polars>=1.39.3",
    "pyqtgraph>=0.14.0",
    "pyside6>=6.11.0",
    "scikit-image>=0.26.0",
    "scipy>=1.17.1",
]
```
