## Supplementary material for "Characterization of Rhizosphere Oxidation Associated with Root Development in Rice Using Planar Oxygen Optodes": RG2DO-Root: README.md

# RG2DO-Root

**RG2DO-Root** is a Python-based analysis tool designed to measure time-series changes in dissolved oxygen concentrations near the rhizosphere using aligned optode images and CT scan slice data.

## Features
* Trace root structures manually on CT and Optode overlay images.
* Calculate average oxygen concentration within a specified radius along the root.
* Export time-series spatial data to CSV for academic analysis.

---

## Prerequisites

This project uses [uv](https://github.com/astral-sh/uv) as a Python package manager.

### Environment Setup
If you have `uv` installed, run:
```bash
uv sync
```
If you prefer not to use `uv`, please refer to the `pyproject.toml` file to install the required dependencies manually via `pip`.

---

## Data Preparation

Place your analysis data in the `target_data` folder. The program expects:
1.  **CT Slice Image**: One aligned CT image.
2.  **Optode Images**: Multiple time-series RGB images.

### File Naming Convention
Files must follow this format (Supported extension: `.png` only):
`[SubjectName]_[Type]_[Note].png`

* **SubjectName**: Name of the subject (e.g., Line/Cultivar name).
* **Type**: Use `CT` for CT slices and `rgb` for optode images.
* **Note**: Information to identify the time point (e.g., date of imaging).

*The program will automatically load these files upon startup based on these keywords.*

---

## Usage

### 1. Launching the Program
Run the following command:
```bash
uv run __main__.py
```

### 2. Interface & Controls
* **Visualization**: The CT slice is displayed by default. Check the **"Oxygen"** box to overlay the latest optode image.
* **Color Scale**: Adjust the oxygen concentration color scale using the slider on the left.
* **Zoom**: Use the mouse wheel to zoom in/out.

### 3. Root Tracing
1.  **Start a Root**: Left-click the base (proximal part) of the root. A red marker and ID `0` will appear.
2.  **Trace**: Move the mouse along the root and **Ctrl + Left-click** to add subsequent points. The points will be connected by lines. Repeat this until you reach the root tip.
3.  **Delete Points**: **Shift + Left-click** to remove the last added point.
4.  **Multiple Roots**: To start a new root (ID `1`, `2`, etc.), simply Left-click the base of the next root and repeat the tracing process.

### 4. Save/Load Progress
* Click **"Save"** in the menu to save your current tracing progress as `target_data.pkl`.
* Click **"Load"** to resume from a previously saved `.pkl` file.

---

## Data Export & Analysis

### Exporting Results
Click the **"Export"** menu to generate CSV files containing oxygen concentration data along the traced roots.

* **Radius Setting**: You can adjust the **Radius** value before exporting. The program calculates the average oxygen concentration of all pixels within the specified radius from the root centerline to reduce noise.
* **Custom Calculation**: To modify the formula for oxygen concentration, edit the `rgb2o` function in `modules/gui.py`.

### Output CSV Format
The exported CSV includes the following columns:

| Column | Description |
| :--- | :--- |
| `root_id` | Unique ID of the root (starting from 0). |
| `distance` | Distance from the root base (pixels). |
| `y` | Y-coordinate. |
| `x` | X-coordinate. |
| `o2` | Calculated oxygen concentration. |
| `count_all` | Total number of pixels within the specified radius. |
| `count_gt1` | Number of pixels with oxygen concentration >= 1%. |
| `suffix` | Metadata extracted from the "Note" section of the filename. |

---

## License

* This program is for personal use or use within the organization to which the user belongs, solely for academic research purposes.
* Any modification of this program, except when necessary for personal or organizational use, requires contacting the copyright holder.
* Commercial use or distribution of this program requires contacting the copyright holder.
* The copyright holder does not guarantee that this program will be free from defects, errors, or malfunctions.
* The copyright holder is not obligated to correct or repair any defects found in the program.
* The copyright holder accepts no responsibility for any direct or indirect damages arising from the use or inability to use this program.
* Users of this program are deemed to be aware that the National Agriculture and Food Research Organization (hereinafter referred to as "NARO") has applied for intellectual property rights such as patents related to the contents of this program.
* NARO will not exercise its intellectual property rights, including patents held by NARO, against users of the program when the program is used solely by those users themselves.
