## Supplementary figures and images for "Characterization of Rhizosphere Oxidation Associated with Root Development in Rice Using Planar Oxygen Optodes"

### test_CT_1230.png

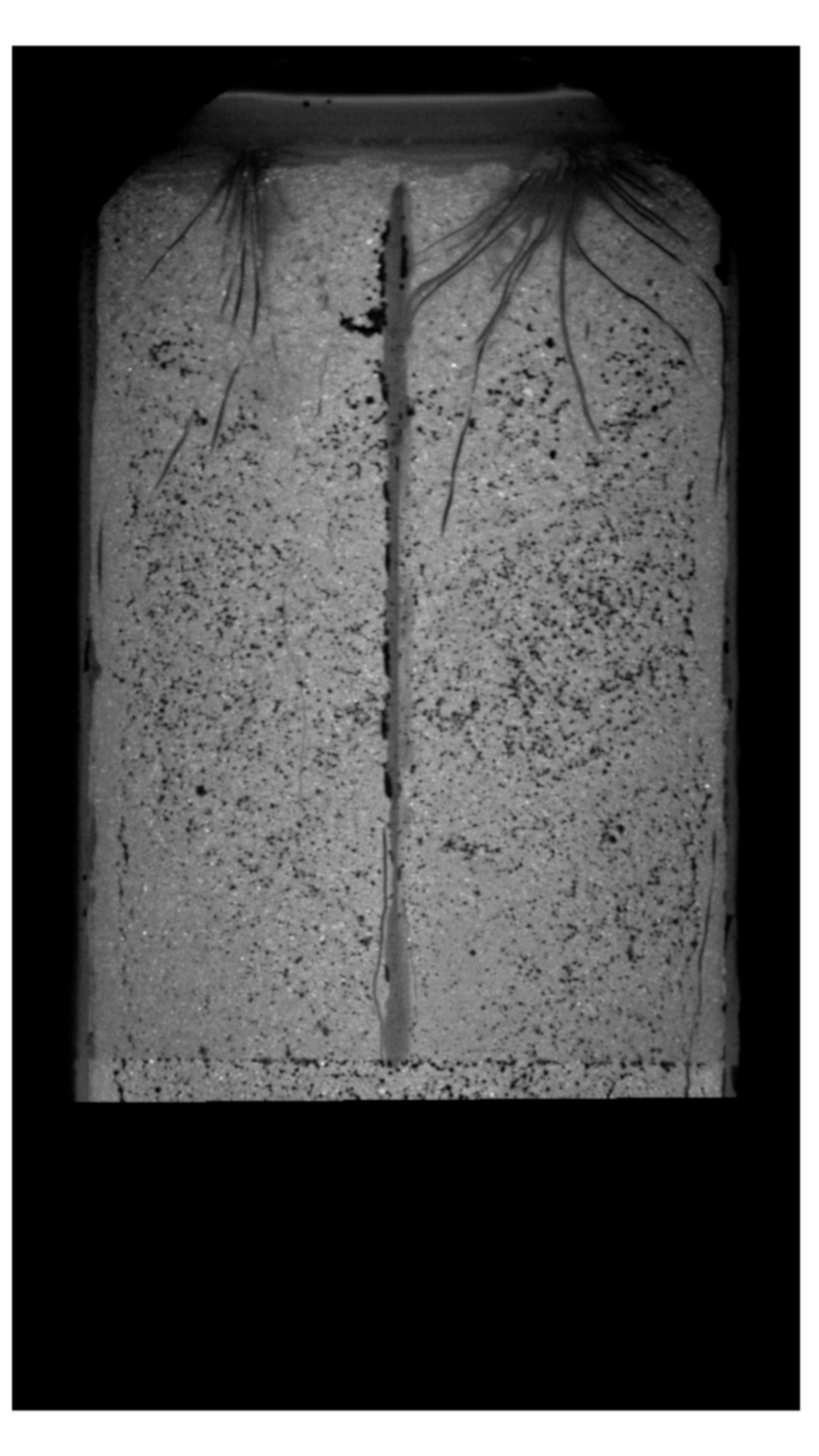

### test_rgb_1220.png

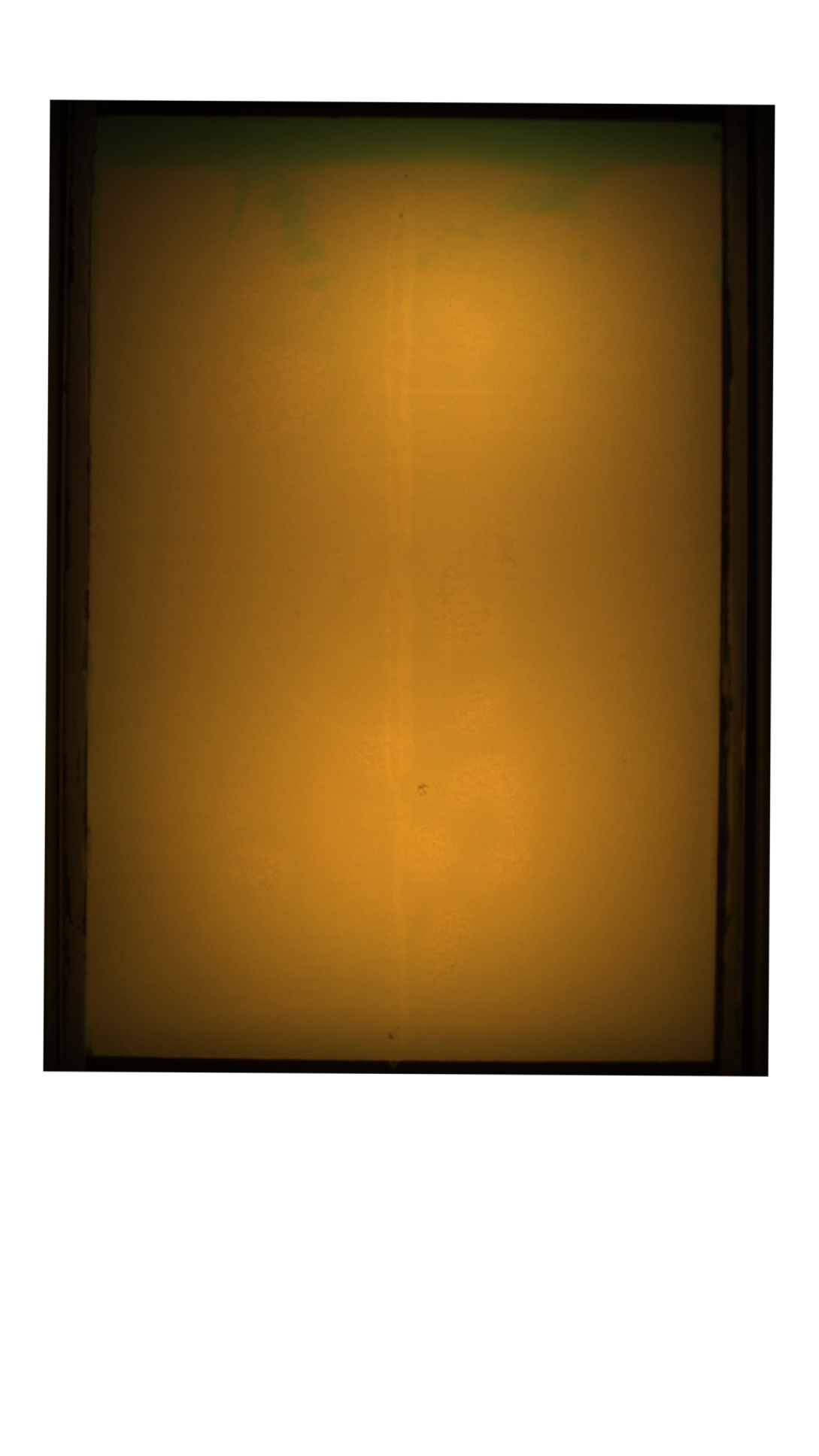

### test_rgb_1225.png

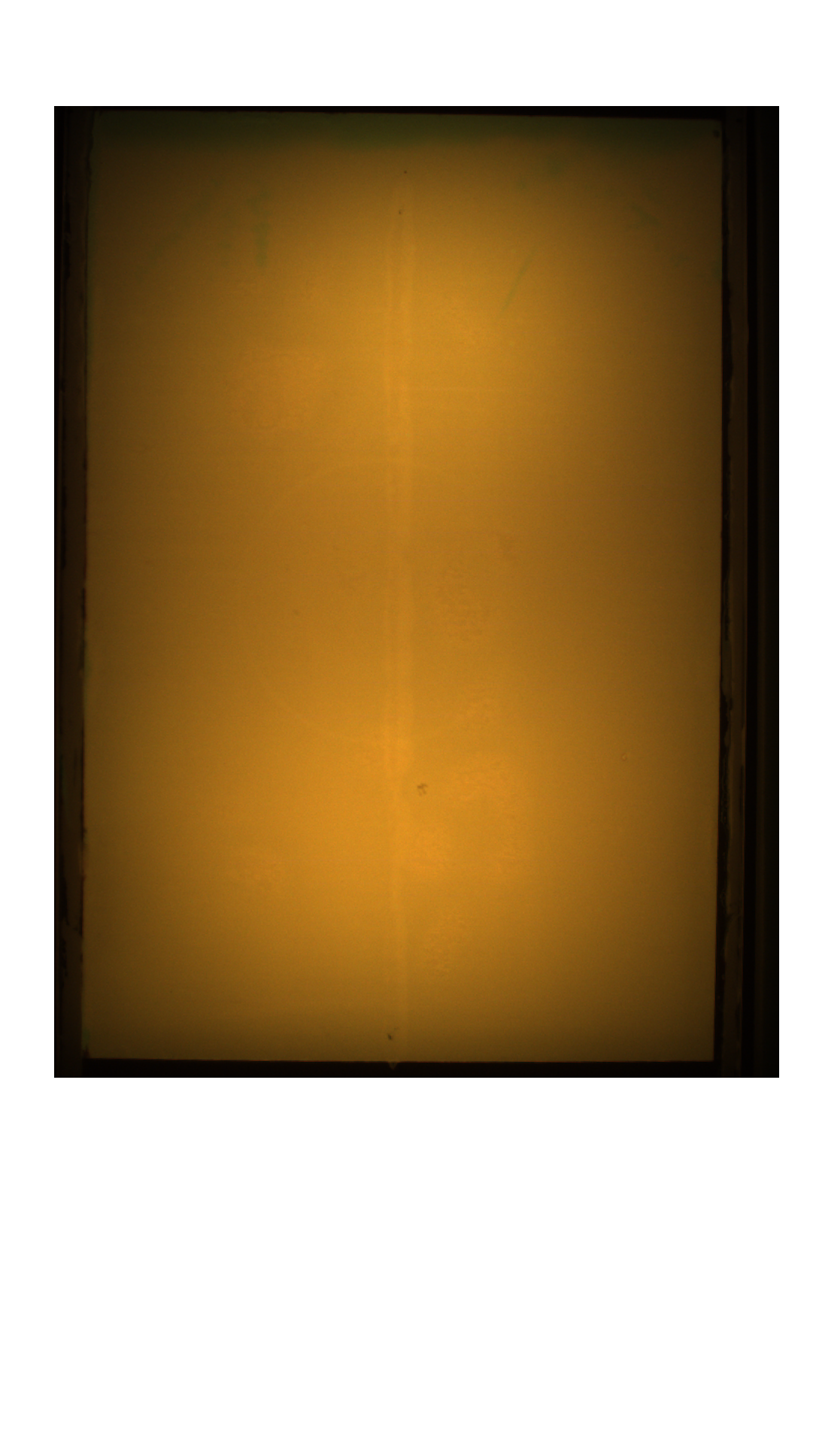

### test_rgb_1230.png

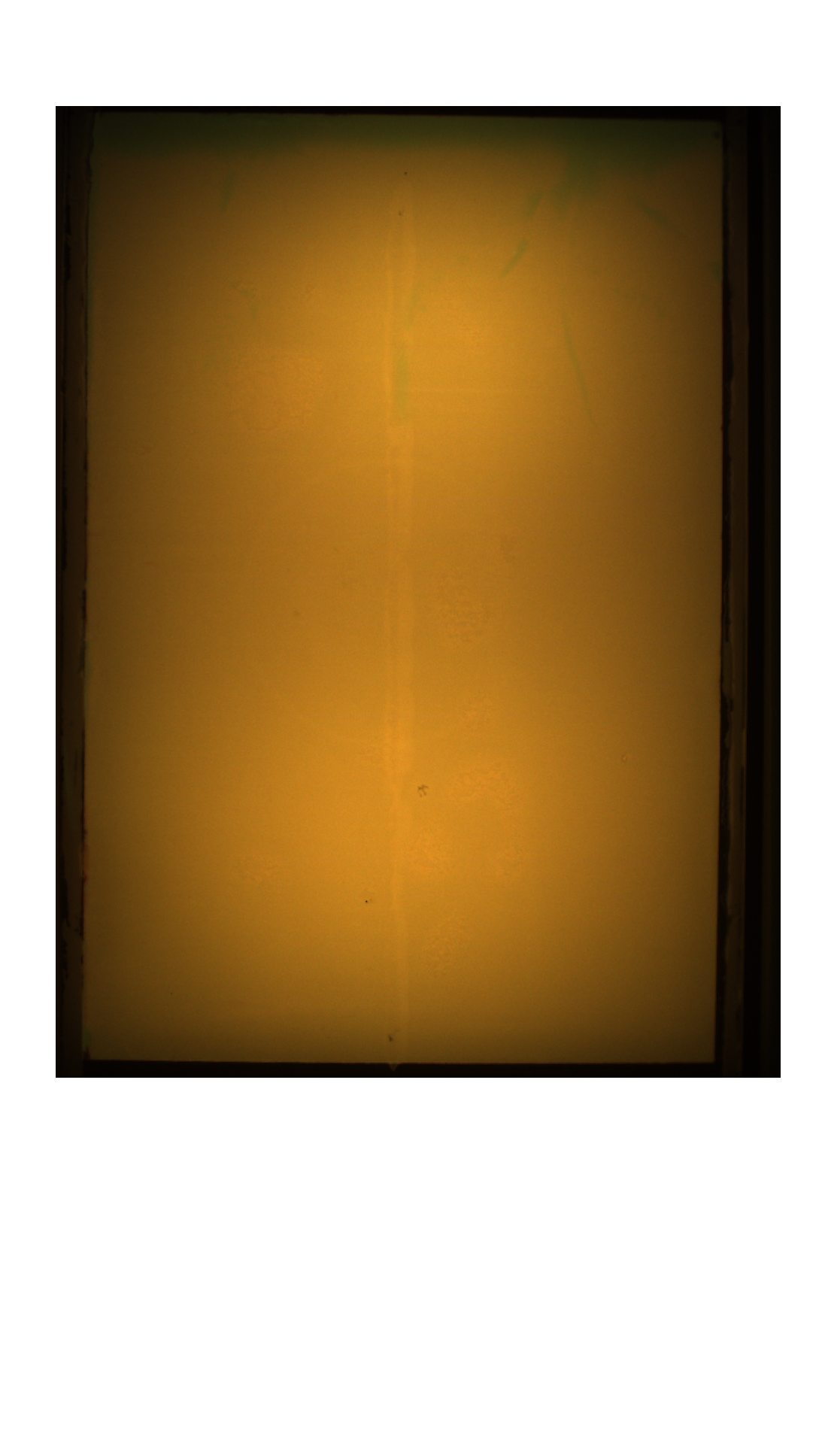
